# Metagenomics Strain Resolution on Assembly Graphs

**DOI:** 10.1101/2020.09.06.284828

**Authors:** Christopher Quince, Sergey Nurk, Sebastien Raguideau, Robert James, Orkun S. Soyer, J. Kimberly Summers, Antoine Limasset, A. Murat Eren, Rayan Chikhi, Aaron E. Darling

**Affiliations:** Warwick Medical School, University of Warwick, Gibbet Hill Road, Coventry, CV4 7AL, UK; Genome Informatics Section, Computational and Statistical Genomics Branch, National Human Genome Research Institute, National Institutes of Health, Bethesda, MD, 20892, USA; School of Life Sciences, University of Warwick, Gibbet Hill Road, Coventry, CV4 7AL, UK; Univ. Lille, CNRS, Inria, UMR 9189 - CRIStAL, France; Department of Medicine, University of Chicago, Chicago, Illinois, USA; Josephine Bay Paul Center, Marine Biological Laboratory, Woods Hole, Massachusetts, USA; Department of Computational Biology, Institut Pasteur, C3BI USR 3756 IP CNRS, Paris, France; The ithree institute, University of Technology Sydney, 15 Broadway, Ultimo, 2007, NSW, Australia

**Keywords:** microbiome, metagenome, strains, Bayesian, microbial community, assembly graph

## Abstract

We introduce a novel bioinformatics pipeline, STrain Resolution ON assembly Graphs (STRONG), which identifies strains *de novo*, when multiple metagenome samples from the same community are available. STRONG performs coassembly, followed by binning into metagenome assembled genomes (MAGs), but uniquely it stores the coassembly graph prior to simplification of variants. This enables the subgraphs for individual single-copy core genes (SCGs) in each MAG to be extracted. It can then thread back reads from the samples to compute per sample coverages for the unitigs in these graphs. These graphs and their unitig coverages are then used in a Bayesian algorithm, BayesPaths, that determines the number of strains present, their sequences or haplotypes on the SCGs and their abundances in each of the samples.

Our approach both avoids the ambiguities of read mapping and allows more of the information on co-occurrence of variants in reads to be utilised than if variants were treated independently, whilst at the same time exploiting the correlation of variants across samples that occurs when they are linked in the same strain. We compare STRONG to the current state of the art on synthetic communities and demonstrate that we can recover more strains, more accurately, and with a realistic estimate of uncertainty deriving from the variational Bayesian algorithm employed for the strain resolution. On a real anaerobic digestor time series we obtained strain-resolved SCGs for over 300 MAGs that for abundant community members match those observed from long Nanopore reads.

## Introduction

There is a growing realisation that to fully understand microbial communities it is necessary to resolve them to the level of individual strains [35]. The strain is for many species the fundamental unit of microbiological diversity. This is because two strains of the same species can have very different functional roles. The classic example is *E. coli*, where one strain can be a dangerous pathogen and another a harmless commensal [24]. The best definition of a strain, and the only one that avoids ambiguity, is a set of clonal descendants of a single cell [15, 39], but strain genomes by this definition can only reliably be determined by sequencing cultured isolates or single cells [30]. The former is not representative of the community and the latter is still too expensive and low-throughput for many applications as well as producing only fragmentary genomes. For these reasons, there is a practical need for efficient methods that can profile microbial communities at high genomic resolution.

In contrast to 16S rRNA gene sequencing, shotgun metagenomics has the potential to resolve microbial communities to the strain level. This is because it generates reads from throughout the genomes of all the community members. It also has the additional advantages of reduced levels of bias and the capability to reconstruct genomes. There are many methods for reference-based strain resolution from metagenome data [1, 35, 42], but they are, and will continue to be, limited by the challenge of comprehensively isolating and sequencing the genomes of diverse microbial strains. Comprehensive reference genome databases may be possible for a few slowly evolving species or particularly well studied pathogens but for the entirety of a complex community it is unlikely to ever be tractable. For example, in a recent *de novo* large-scale binning study of the relatively well-studied human gut microbiome, it was found that 77% of the species recovered did not have a reference genome in public databases [31]. This suggests that even less of the strain-level diversity in those samples would be represented in a genome database. These observations motivate the need for *de novo* methods of metagenomic strain resolution.

In the metagenomics context, we adopt the definition of a ‘metagenome strain’ as a clonal subpopulation with sufficiently low levels of recombination with other strains, that it can be distinguished genetically from them. This does not require that recombination between strains does not occur, rather that either because of physical separation or selection, it has not been sufficiently strong relative to the rate of mutation [40], to generate a continuum of diversity throughout the genome. This means members of a ‘metagenome strain’ may differ substantially from each other particularly in rapidly evolving accessory regions and the subpopulation as a whole may descend from multiple cells but with a core genome that has descended from a single cell in the recent past. This is equivalent to the definition of ‘lineage’ in [29]. For ease, in the discussion below we will refer to strain in the metagenome context when properly we mean this looser definition of a strain as a genetically distinct subpopulation.

*De novo* assembly of genomes from short read metagenome sequences remains very challenging. Assemblies become fragmented for two reasons: firstly, low coverage genomes will fragment through chance occurrences where sequence coverage drops out, following Lander and Waterman statistics [17], secondly, if either intra or inter-genomic repeats are present then the assembly graphs used to represent possible sequence overlaps become very complex, and it is unclear which paths correspond to true genomes. Both of these issues are particularly problematic for metagenomes, where there can be a wide range of species abundances, and in a complex community a significant fraction of the species may be at low coverage. The first challenge can be addressed by sequencing more deeply. More difficult to address is the problem of repeats. Just as they do in isolate genome sequencing, intra-genomic repeats such as the 16S rRNA operon will lead to uncertainty in metagenomic assemblies, but if multiple closely related strains from the same species are present then they will possess potentially large regions of shared sequence. If the strain genomes are of comparable divergence to the reciprocal of the read length then very complex graphs will result, for typical short read sequencing (75-150bp) this would be strains at around 98-99.5% sequence identity. The result is that it is not possible to find long paths in the graph that can be unambiguously assembled into long contiguous sequence or contigs. For this reason metagenome assemblies for strain-diverse communities can comprise millions of contigs when made from short read data, with the added drawback that in the metagenomics context, we do not even know which contig derives from which species. For species that contain multiple very similar strains (*>* 99.9%), then we expect better assemblies but the variants are then too far apart to be linked or phased by Illumina reads. In that case we may resolve the large-scale genome structure but not the sequences of the individual strains, which we will refer to as their haplotypes.

Metagenomic contig binning methods attempt to mitigate the problem introduced by standard metagenome sample processing approaches, wherein the origin of each sequence read is unknown. Contig binning works because contigs deriving from the same or similar genomes will share features that can be learnt without prior knowledge. These features can be sequence composition, but it is also possible to use per-sample coverage depths of contigs as a more powerful feature, if multiple samples are available from the same (or very similar) communities [2]. There are now numerous algorithms capable of using both coverage across samples and composition to automatically cluster contigs and determine from single-copy core gene (SCG) frequencies where the resulting bins are good quality metagenome assembled genomes (MAGs) [3, 13]. These tools enable genome bins to be extracted *de novo* from metagenomes, and are becoming crucial for studying unculturable organisms, contributing to many exciting discoveries, such as the description of the Candidate Phyla Radiation [9] or an improved understanding of the diversity of nitrogen fixers in the open ocean [14].

The resolution of genome binning though, is limited by the resolution of the assembler, with a typical maximum kmer length of around 100, the best case is that we can resolve to about 1% sequence divergence, so that bins correspond to something between a species and a strain. In the presence of strain diversity, those contigs that are shared across strains will become a consensus of the strains present, in the ideal situation their sequence would be that of the most abundant strain, but even this is not guaranteed. Contigs that are part of the accessory genome and present in a subset of strains may be successfully binned with the core genome, but they may not if they are too short or divergent in coverage. Consequently, if multiple strains are present in the assembly the MAGs that result from binning will be an imperfect composite of multiple strains.

Strains in a metagenome can exhibit variation in shared genes, such as insertions/deletions and single-nucleotide variants or SNVs, as well as in their accessory gene complements. Recently, we introduced DESMAN [32] to resolve subpopulations in MAGs using variant frequencies on contigs when multiple samples from a community are available. This is similar to contig binning using coverage but it can be viewed as a relaxed form of clustering closer to non-negative matrix factorisation, because each variant can appear in more than one subpopulation haplotype. Similar strategies had been proposed prior to DESMAN but using variant frequencies on reference genomes e.g. Lineages [29] and Constrains [27]. DESMAN and other earlier methods are all ‘linear mapping-based methods’ where metagenomic reads are mapped onto a linear sequence, either a reference or consensus contig. This has multiple drawbacks: firstly, the type of variant that can be represented is limited to changes at a single base; secondly, mapping onto a linear sequence can be challenging when there is variation present yielding unreliable results [19]; thirdly, it treats every variant as independent ignoring the co-occurrence of variants in reads, which is a powerful extra source of information when strain divergence is greater than the inverse of read length, when we would expect most reads to contain more than one variant. The last issue can be addressed by keeping track of which variants appear in which reads but that requires extra bookkeeping [18].

To address these limitations, we introduce a new method, STRONG (Strain Resolution ON Graphs), for analysing metagenome series when multiple samples are available either from the same microbial community e.g. longitudinal time-series or cross-sectional studies where the communities are similar enough to share a significant fraction of strains. STRONG can determine the number of ‘metagenome strains’ in a MAG formed from binning of a coassembly of all the samples, together with their sequences across multiple single-copy core genes, which we define as the strain haplotype, and the coverages of each strain in each sample. STRONG avoids the limitations of the variant-based approaches by resolving haplotypes directly on assembly graphs using a novel variational Bayesian algorithm, BayesPaths.

This graph-based approach allows more complex variant structure and incorporates read information. The usefulness of graphs for understanding microbial strains has been noted before, and efficient algorithms developed for querying complex graphs and extracting more complete representatives of MAGs in the presence of strain diversity [10]. STRONG, however, is the first time that graphs have been used in an automated workflow to actually decompose that strain diversity into haplotypes across multiple genes using multiple samples. We compare STRONG to the current state of the art, DESMAN, on synthetic microbial communities and a real metagenome time series from an anaerobic digester. In the former case we validate using the known genome sequences, and for the latter we compare abundant MAGs with haplotypes derived independently from Oxford Nanopore MinION long reads.

## Results

### STRONG pipeline

The detailed pipeline is described in the Methods but the key steps are summarised in Figure 1 and reiterated here. We start from multiple samples of the same community and jointly coassemble them with metaSPAdes, we save a high resolution graph (HRG) early in the assembly process that preserves all the variant information in the coassembly. The metaSPAdes assembly process then proceeds as normal and the resulting contigs are binned using CONCOCT. We annotate the single-copy core genes in the contigs, allowing us to identify a subset of bins as MAGs. A novel algorithm was then developed to map these SCG ORFs onto the HRG and extract the complete assembly subgraphs corresponding to the genes of interest (Methods - Relevant subgraph extraction). We obtained per sample unitig coverages on these subgraphs by threading reads directly onto them. These subgraphs were simplified with a noise filtering algorithm that used the MAG coverage depths, calculated as the length weighted average of the contigs assigned to that MAG. The simplified subgraphs contain all the information required for the BayesPaths algorithm (Methods - BayesPaths), that simultaneously solves for the number of strains present, their coverage in each sample, and their sequences on the SCGs. SCGs from the same MAG are linked through the binning process and jointly solved in the strain resolution procedure to generate linked strain resolved sequences for each SCG. We will refer below to the SCG sequences for a given strain as its haplotype. The pipeline also applies DESMAN [32], to the same MAGs for comparative purposes, and will perform benchmarking if known genomes are available. It is important to note that some SCGs will be filtered during the BayesPaths procedure, see Methods, so that sequence inference is only performed on a subset in the final output.

**Figure 1.**
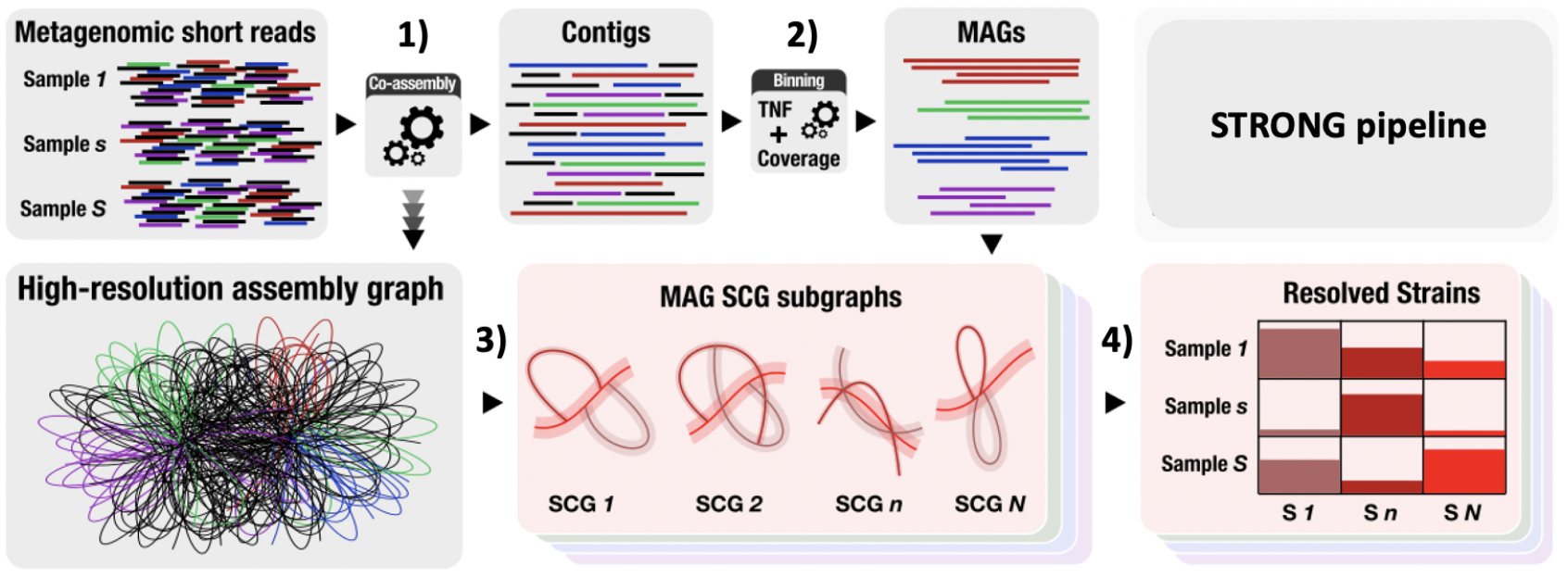
STRONG pipeline. This figure illustrates the principal steps in the STRONG pipeline (see Methods - STRONG Pipeline). Step 1) Co-assembly with metaSPAdes and storage of a high-resolution graph (HRG). Step 2) Contig binning with CONCOCT and annotation of single-copy core genes (SCGs). Step 3) Mapping of SCGs onto the HRG and extraction of individual SCG assembly graphs together with per-sample unitig coverages. Step 4) Joint solution of SCG assembly graphs from each MAG with BayesPaths to determine strain number, haplotypes and per-sample coverages.

### Synthetic data sets

In order to provide an example metagenome data set with a known strain configuration for each species, we created a synthetic community comprised of 100 strains, with known genomes deriving from 45 species, with 20 species represented by a single strain, 10 with two strains, 5 with three, 5 with four and 5 species with five strains. We then generated four data sets from this community with the same total number of reads (150 million 2×150 bp) but increasing sample numbers (3, 5, 10 and 15 samples). This configuration, where most species have a single strain, might be an appropriate approximation to the human gut microbiome [38]. We denote these data sets Synth S03, Synth S05, Synth S10 and Synth S15. For each sample number, random species abundances were generated from a log-normal distribution, with strain proportions from a Dirichlet. Full details of the synthetic sequence generation are given in the Methods.

The STRONG pipeline was then applied to each of these data sets in turn. In Figure 2 we illustrate the STRONG output for a single gene, COG0532 ‘Translation initiation factor IF-2’ [37], from one MAG, Bin 55 of the ten sample synthetic data set, giving the resulting decomposition of the assembly subgraph into three strains. Noting that the strains were resolved in this MAG over 22 single-copy core genes simultaneously, and that for this 3.4 kbp gene the haplotypes were found without errors.

**Figure 2.**
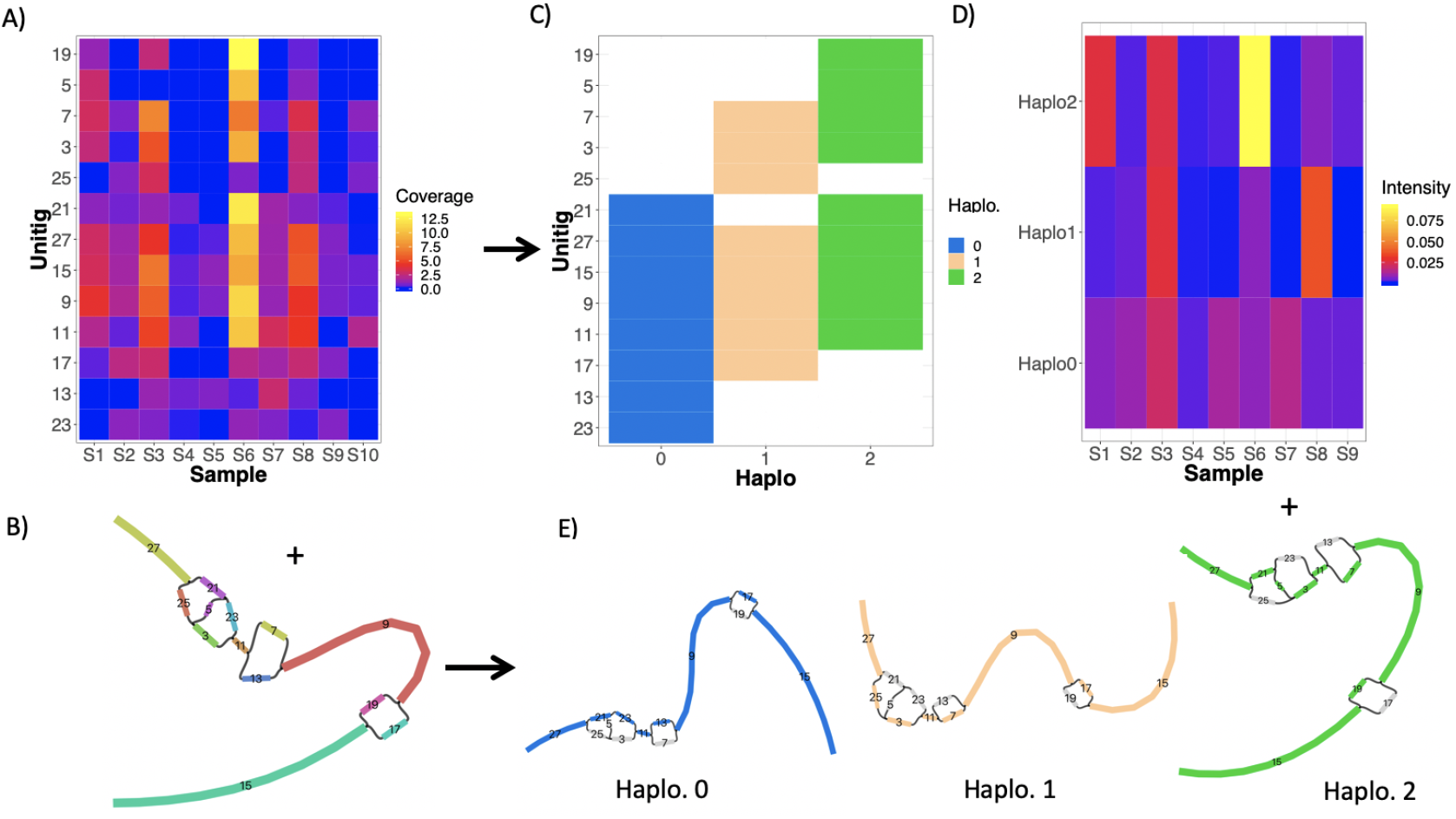
BayesPaths algorithm. This illustrates the BayesPaths algorithm for a single COG0532 from one MAG, Bin 55 of the ten sample synthetic data set. The algorithm predicted 3 strains. We show the input to the algorithm: A) the unitig coverages across samples plus B) the unitig graph without strain assignments. The outputs of the algorithm are shown in C) the assignments of haplotypes to each unitig, D) the strain intensities across samples, effectively coverage divided by read length (see Methods - BayesPaths), and E) unitig graphs for each haplotype with their most likely paths. This algorithm is explained in detail in the Methods - BayesPaths.

For each of the four synthetic data sets we considered only MAGs which were assigned to species (see Methods) with at least two strains - 20, 21, 24 and 22 MAGs, from the Synth S03, Synth S05, Synth S10 and Synth S15 data sets respectively. For each MAG we mapped the predicted haplotypes for the optimal strain decomposition for both the STRONG pipeline and DESMAN algorithms onto the known reference strains. We then assigned each haplotype prediction to its best matching reference. The best such match was denoted ‘Found’. If multiple predicted haplotypes matched to the same reference they were denoted as ‘Repeated’. If a reference had no haplotype prediction that matched to it better than the other references, it was denoted as ‘Not found’. For the aggregate across these MAGs we show the total number of such strains for each of the four data sets in Figure 3.

**Figure 3.**
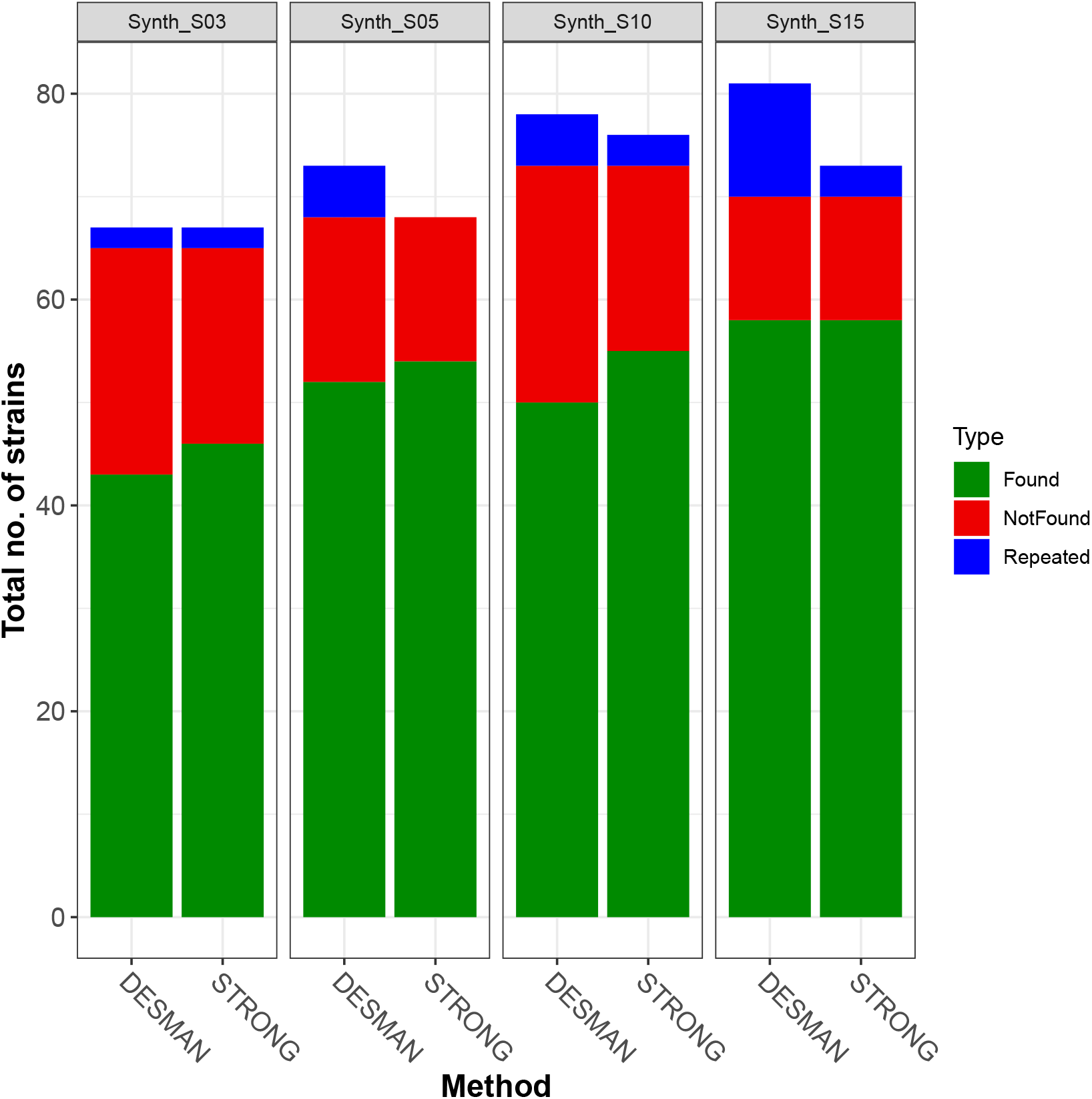
No. of strains resolved by STRONG and DESMAN algorithms in the synthetic community data sets. For MAGs with two or more strains we mapped haplotypes to the references and assigned each predicted haplotype to its best matching reference. The best such match was denoted ‘Found’. If multiple haplotypes matched to the same reference they were denoted as ‘Repeated’. If a reference had no predicted haplotypes matched to it, it was denoted as ‘Not found’. The bars give the total numbers in each category summed over MAGs for the two methods (DESMAN and STRONG) and the panels results for the four different data sets with increasing number of samples (Synth_S03, Synth_S05, Synth_S10 and Synth_S15).

STRONG consistently outperforms DESMAN in terms of number of strains found, in total across all four samples it resolved 213 strains vs. 200 for DESMAN *i*.*e*. a 6.5% increase. It also had fewer ‘Repeated’ strains, 8 vs. 23: a reduction of 65%. The strains ‘Found’ were also reconstructed more accurately, the per base error rate for the BayesPaths reconstructions averaged across all MAGs and all data sets was just 0.052%, three times lower than that for DESMAN, 0.176%. This improvement was observed for all four data sets (see Table 1 and Figure 4). STRONG was more likely to predict the correct number of strains, doing so for 73% of MAGs summed across samples numbers versus 60% for DESMAN. It was also better at predicting the strain relative abundances. Regressing true abundance against predicted abundance gave an adjusted *R*^2^ of 0.84 averaged across sample numbers for STRONG vs. 0.80 for DESMAN. When this was restricted to MAGs where the number of strains was correctly predicted, then both algorithms did better but STRONG still out performed DESMAN, with a mean *R*^2^ of 0.98 compared to 0.93. Although the quantity varied across the four data sets, roughly 1/3 of the SCGs were filtered during the BayesPaths as outliers (see - Table 1).

**Table 1.** Comparison of STRONG to DESMAN for strain reconstruction in the synthetic community data sets. Data set: Results are shown for the four different sample numbers. MAGs: The number of MAGs reconstructed with more than two reference strains. #SCGs: The average number of SCGs found in each MAG. #fSCGs The average number of SCGs after filtering in STRONG. Found: Number of reference strains that had a predicted strain that best matched it. Not F.: Number of reference strains that had no predicted strain with a closest match to it. Rep.: Number of reference strains with more than one best matching predicted strain. Err: The average error rate of the ‘Found’ strains in percentage base pairs. *R*^2^: Correlation between predicted and actual strain relative proportions given as adjusted *R*^2^, the figure in parentheses is when restricted to MAGs where the number of strains was correctly predicted. *f* ^*G*^: the fraction of MAGs where the number of strains was correctly inferred.

| Method | Data set | MAGs | #SCGs | #fSCGs | Found | Not F. | Rep. | Err | $R^2$ | $f^G$ |
| --- | --- | --- | --- | --- | --- | --- | --- | --- | --- | --- |
| STRONG<br>DESMAN | Synth_S03 | 20 | 35.75 | 18.95 | 46<br>43 | 19<br>22 | 2<br>2 | 0.068<br>0.121 | 0.86 (0.99)<br>0.79 (0.93) | 23/34 = 0.68<br>24/34 = 0.71 |
| STRONG<br>DESMAN | Synth_S05 | 21 | 35.29 | 22.14 | 54<br>52 | 14<br>16 | 0<br>5 | 0.054<br>0.186 | 0.82 (0.98)<br>0.83 (0.99) | 28/37 = 0.76<br>20/37 = 0.54 |
| STRONG<br>DESMAN | Synth_S10 | 24 | 32.23 | 21.91 | 55<br>50 | 18<br>23 | 3<br>5 | 0.042<br>0.252 | 0.83 (0.99)<br>0.76 (0.95) | 26/39 = 0.67<br>21/39 = 0.54 |
| STRONG<br>DESMAN | Synth_S15 | 22 | 35.45 | 23.36 | 58<br>58 | 12<br>12 | 3<br>11 | 0.045<br>0.143 | 0.86 (0.98)<br>0.81 (0.87) | 32/40 = 0.80<br>25/40 = 0.63 |

**Figure 4.**
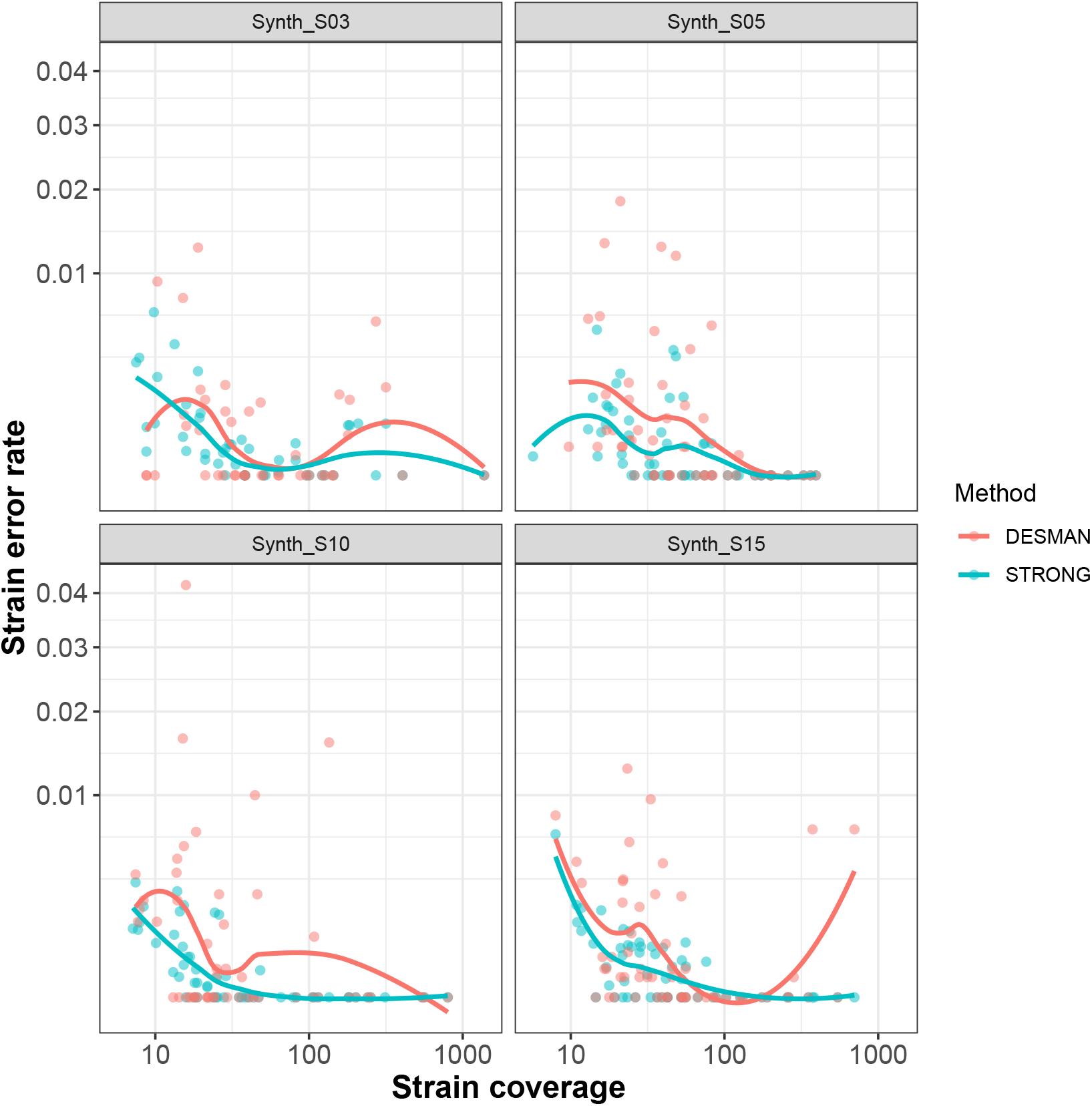
Error rates in strains found against coverage depth for STRONG and DESMAN algorithms in the synthetic community data sets. For the ‘Found’ strains we computed per base error rate to the matched reference, this is shown on the y-axis, against strain total coverage depth summed across samples on the x-axis, both axes are log transformed. The results are separated across methods (DESMAN and STRONG) and sample number in the synthetic community.

The STRONG pipeline outperforms DESMAN, but it still misses strains that are present. In total across all MAGs and data sets, 63/276 *i*.*e*. 22.8%, of strains were missed by STRONG. Some of these, 7 out of 63, were below the minimum coverage of detected strains (5.68), but most were not, suggesting that either they were not sufficiently divergent in terms of nucleotides or coverage profiles to be detected. Examination of phylogenetic trees for the haplotypes and reference genomes constructed using the SCGs revealed that in many cases ‘Not found’ strains had identical SCG haplotypes to those that were resolved.

The BayesPaths algorithm used to resolve strains in STRONG uses variational inference (see Methods - BayesPaths), an approximate Bayesian strategy [7]. This has the advantage of providing estimates of uncertainty in the inference of both the strain haplotypes and their abundances. The algorithm predicts the marginal probabilities that a given strain passes through a particular unitig. To provide a single sequence for the evaluation above and applications below we output the most likely path and hence sequence for each strain. However, we also calculate an estimate of path uncertainty by sampling many possible paths (default 100) consistent with the marginal distributions and calculate the average number of nodes that deviate from the most likely path, we refer to this as the divergence. For the ‘Found’ strains this correlates strongly with actual error rate to the reference strain (Pearson’s correlation *r* = 0.56, *p* < 2.2*e −* 16 - see Figure S1). Thus the divergence is a useful prediction of uncertainty in the haplotype sequence inference, enabling us to estimate error rates in real data sets in the absence of known reference sequences. Roughly speaking, the expected per base error rate is 0.01 times the divergence, so that a strain divergence of 0.1 predicts a 0.1% error rate. In real data sets, the uncertainty estimates in the abundances are also useful, placing bounds on the abundance of individual strains in each sample.

In Table S3 we give approximate run times for each component of the STRONG pipeline on the synthetic community data sets, using 64 threads on a standard bioinformatics server (see Table S3). The BayesPaths step is the most time consuming part of the analysis (up to 36 hours), but it is still comparable to the initial coassembly. The only part of the pipeline with substantial memory requirements is the initial coassembly with metaSPAdes, the other steps are CPU limited.

### Anaerobic digester time series

We next applied the STRONG pipeline to a real metagenomics time series, comprising ten samples taken at approximately 5 weekly intervals, from an industrial anaerobic digestion reactor (see Table S4 and Methods for details). This provides an evaluation community of intermediate complexity to test the pipeline’s capability to resolve strains and reconstruct intraspecies dynamics. Each sample was sequenced on the NovaSeq platform with 2×150 bp reads at a mean depth of 11.63 Gbp. One sample was also run on a Nanopore MinION flow cell producing 43.78 Gbp of reads with a read N50 of 6,727 bp and a maximum length of 108 kbp.

CONCOCT binning produced 905 bins, of which 309 had 75% of SCGs present in single-copy, which we designate MAGs. In total 11 of these MAGs exhibited overlapping SCG graphs and were merged into 6 composite MAGs (see Methods - STRONG Pipeline), so that 304 MAGs were actually used in the strain decomposition. We calculated coverage depth per sample for each bin and then normalised by sample size to obtain a community profile at each time point. Overall the reactor exhibited a clear shift in community structure over time, despite consistent operating conditions, with sample time explaining 48% of the variation in community structure (*p* = 0.001 - Figure S2). Of the MAGs, 110 had an abundance that changed significantly over time (Bonferonni adjusted p-value < 0.05 from Pearson’s correlation of log transformed normalised abundance) and these were evenly split between those that increased (55) or decreased in abundance (55).

We used the STRONG algorithm to resolve strains in the 304 MAGs. This is a complex data set and running the complete pipeline took over 16 days, of which roughly 60% of the time was spent on the BayesPaths strain resolution (see Table S3). The number of strains found varied between 1 and 7, with a mean of 1.7, shown as a function of coverage depth in Figure 5. In total 121 (39.8%) of these MAGs had more than one strain, and there was a significant positive association between MAG coverage depth and number of strains (*r* = 0.36, *p* = 1.004*e −* 10), which is expected, as low coverage MAGs will be under-sampled. This correlation disappears though when we restrict to all MAGs with a coverage greater than thirty (*r* = 0.19, *p* = 0.1023). On average 20.9 SCGs were used after filtering for strain haplotype predictions.

**Figure 5.**
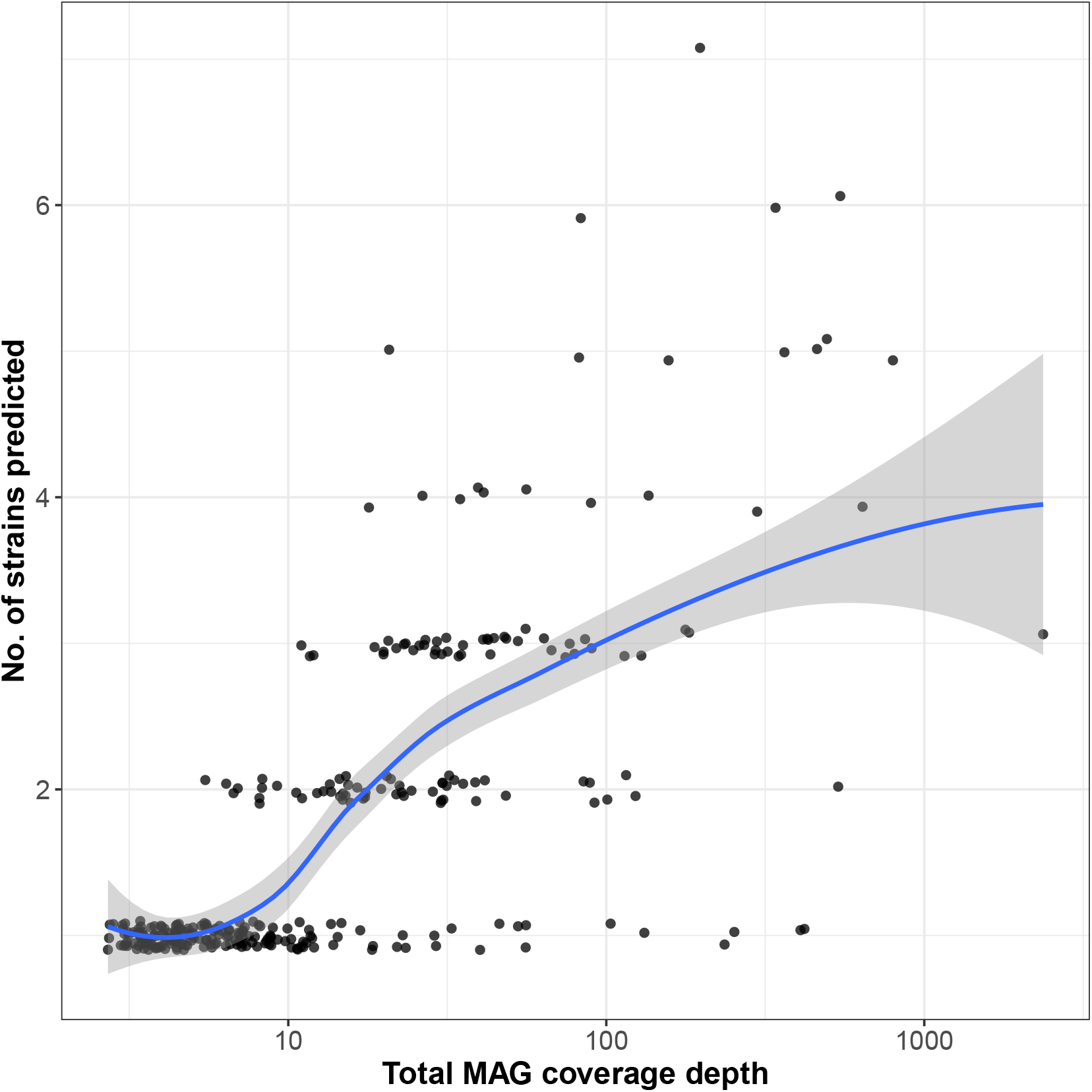
Number of strains resolved by STRONG against MAG coverage depth for the AD time series. Pearson’s correlation between coverage depth and number of strains (*r* = 0.36, *p* = 1.004*e −* 10). The curve indicates a LOESS smoothing.

For the 108 MAGs that had at least two strains with relative frequencies determined in five or more samples we used permutation ANOVA to determine whether strain proportions depended on sampling time. In total 13 of the MAGs had an adjusted p-value < 0.05 *i*.*e*. 12.0%. For these same MAGs 37 had a total coverage that changed significantly over time with an adjusted p-value < 0.05 *i*.*e*. 34.2%. Therefore the intra-species dynamics are more stable than inter-species, with strain proportions remaining fixed as the MAG coverages vary, this was true for 33 of the 37 MAGs that changed significantly in coverage.

In Figure 6, we use the Anvi’o program [16] to summarise information on phylogeny, taxonomy, normalised coverages in the ten samples, and whether the MAGs changed significantly in total abundance, together with the number of strains resolved by STRONG and if those strain relative proportions changed significantly with time. This was restricted to just those 114 MAGs with an aggregate coverage greater than twenty to simplify the diagram.

**Figure 6.**
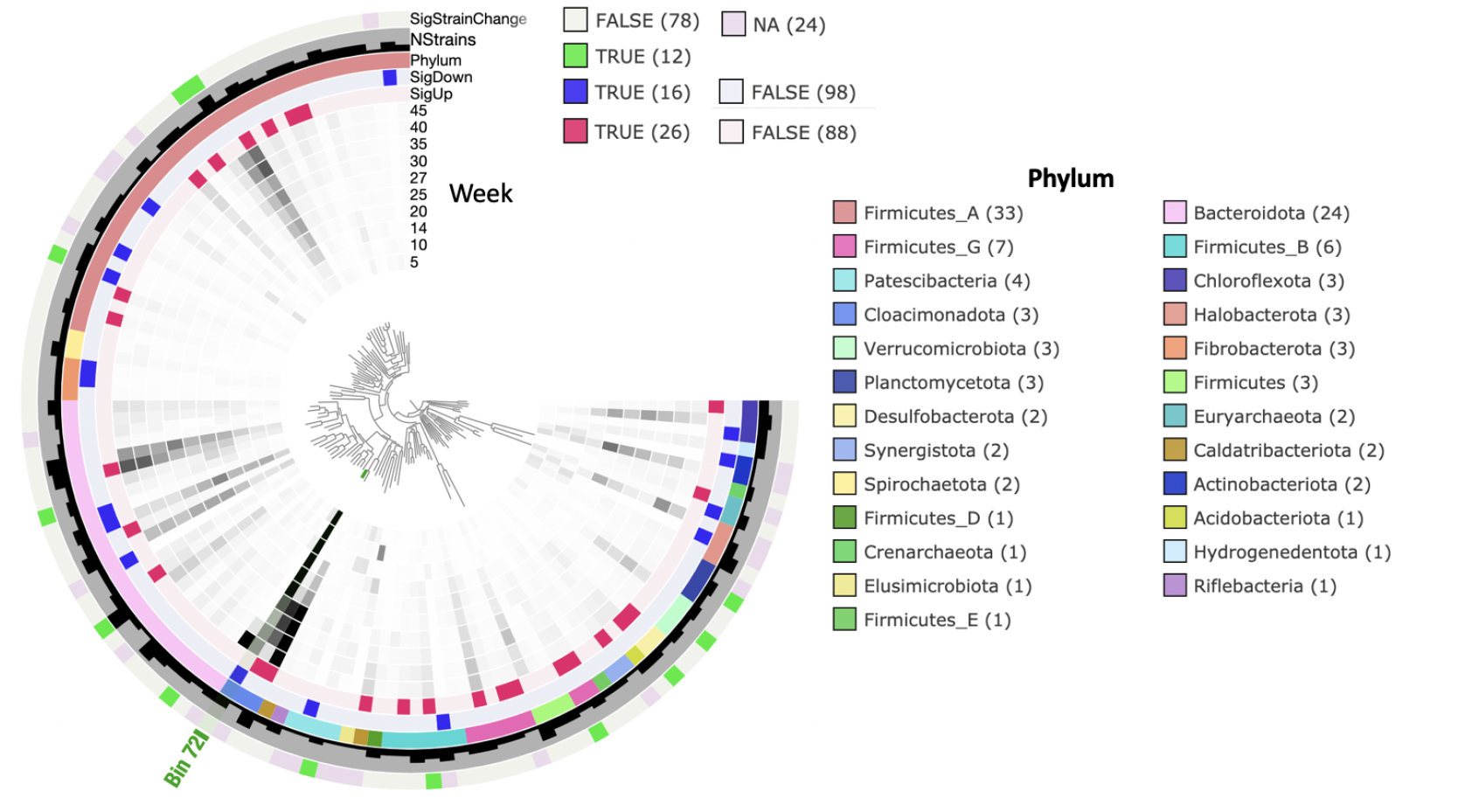
MAG summary for anaerobic digester time series. For the 114 MAGs with aggregate coverage *>* 20 we give their phylogeny constructed using concatenated marker genes together with their normalised coverages in the ten samples. We also indicate which MAGs significantly increased (SigUp) or decreased (SigDown) in total abundance (adjusted *p* < 0.05), their GTDB phylum assignment, no. of strains resolved by STRONG and whether the strain abundances changed significantly over time (adjusted *p* < 0.05) using permutation ANOVA (SigStrainChange).

The Nanopore sequencing provides us with a means to directly test the validity of the STRONG haplotype reconstructions, at least for the most abundant MAGs. The most abundant MAG, Bin_72, had an aggregate short read coverage depth of 2364.25, across all the samples. This MAG was assigned to the phylum Cloacimonadota using the GTDB taxonomy [11]. Interestingly, this is an example of a MAG which changes significantly in abundance, decreasing over time, (adjusted *p* = 4.9*e* 05) but where the proportions of the three strains predicted varied less dramatically (*R*^2^ = 0.35 adjusted *p* = 0.089) - see Figure S5. We will focus on the longest SCG for which strains were resolved, COG0532, where the three strains are present in only two variants, haplotypes 0 and 2 being identical on this core gene. In Figure S4 we give the short read variant graph for this gene, which in this case is mostly simple bubbles, together with the assigned haplotypes. In fact, across the 18 SCGs used to decompose strains, haplotypes 0 and 1 were most similar with 99.7% nucleotide identity. These two strains had 99.4% and 99.1% identity with haplotype 2 respectively. That this pattern was not observed on COG0532 may suggest some recombination in the evolution of these organisms.

In Figure 7 we show for COG0532 both the Nanopore reads that map to this gene and the three haplotypes inferred by BayesPaths, as an Non-metric Multi-dimensional Scaling (NMDS) plot using fractional Hamming distances on the short read variant positions. These are defined as the Hamming distance between two reads but only on the intersecting variant positions and ignoring gaps. We then normalise by the number of such non-gap intersecting positions to give a distance between 0 and 1. The Nanopore reads are consistent with the inference of two variants on this gene, as there are two clear clusters observed, and the two modes of those clusters are close to those haplotypes. The variation around the modes is caused by the high error rate of the Nanopore reads.

**Figure 7.**
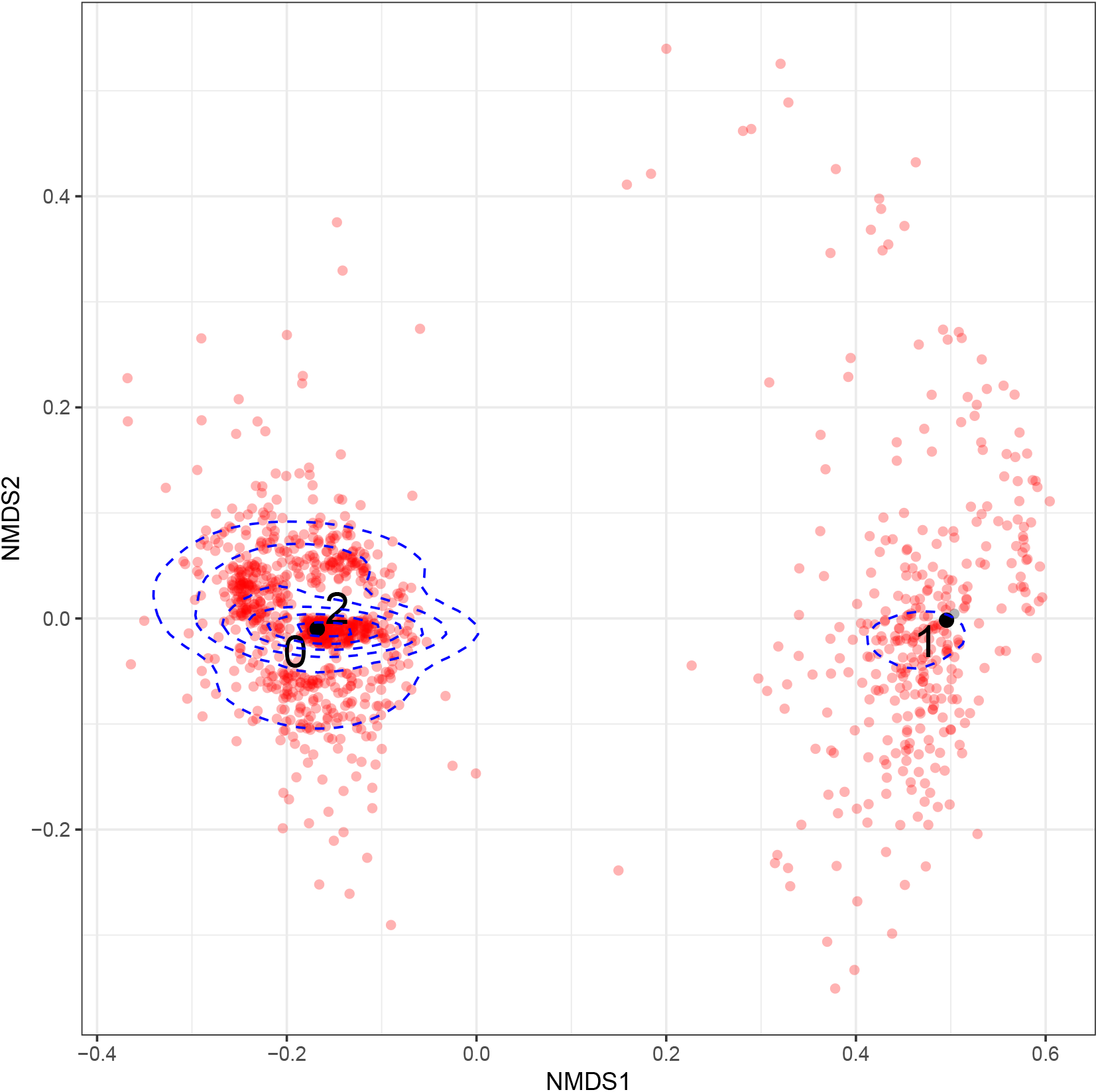
Comparison of Nanopore reads to STRONG prediction for COG0532 from Bin_72. Non-metric multidimensional scaling of Nanopore reads that mapped to COG0532 from Bin_72 of the anaerobic digester time series (red) together with the three haplotypes reconstructed from short reads by STRONG (black 0, 1 and 2). Haplotypes 0 and 2 were identical for COG0532. Distances were calculated as fractional Hamming distances (see text) on short read variant positions (see Methods - Nanopore Sequence Analysis). Blue dashed lines indicate read density contours.

In order to provide a quantitative comparison of the Nanopore reads and the STRONG predictions, we applied the EM algorithm defined in the Methods (Nanopore Sequence Analysis) on the 1,603 Nanopore reads mapping to this COG (cluster of orthologous groups). Examining the negative log-likelihood as a function of number of strains, it flattens at two strains (see Figure S3) and the two strains inferred exactly match (100% identity over 2,313 bps) haplotypes 0/2 and 1 respectively. Furthermore, STRONG in this sample predicted frequencies of 28.0% for haplotype 1. This closely matched the Nanopore haplotype frequencies for this strain of 27.6%. We also ran the Nanopore EM algorithm for all 18 filtered COGs in this bin separately. For the 11 COGs where more than one strain was predicted from the Nanopore reads, we compared the STRONG and Nanopore predictions. For haplotypes 0, 1 and 2 exact matches were found for 6, 7 and 4 SCGs respectively with average nucleotide identities across all genes of 99.89%, 99.89% and 99.82%.

For lower coverage MAGs we generally obtain a reasonable correspondence between the STRONG haplotypes and Nanopore predictions. In most cases the number of strains is comparable between the two, although the accuracy of matches reduces with decreased Nanopore read counts, as we might expect. As an example, in Figure S7 we compare Nanopore reads with the five STRONG haplotypes from COG0072 of Bin 846, a Firmicutes MAG in the AD time series. The most abundant Nanopore mode clearly matches STRONG haplotype 4, the most abundant strain in this sample, and there is also some support for haplotypes 0 and 2. There is less evidence for strains 1 and 3, but these are low abundance in this sample (see Figure S9). This is confirmed from the EM algorithm applied to the Nanopore reads matching this gene, where we would predict four Nanopore haplotypes (Figure S6). Comparing these 4 Nanopore strains to the STRONG predictions we find that three closely match: Nanopore haplotype 0 matched best to STRONG strain 4 with 98.8% nucleotide identity, Nanopore haplotype 1 to STRONG 4 with 99.9% identity and Nanopore haplotype 2 to STRONG strain 0 with 99.7% identity. There is also a correspondence in relative abundance, with the most abundant Nanopore haplotype 1 recruiting 82% of the reads vs 74% relative frequency for the corresponding strain haplotype from STRONG.

## Discussion

We have demonstrated that on synthetic data the STRONG pipeline and the BayesPaths algorithm are able to accurately infer strain sequences on the SCGs and abundances across samples. Performance does improve with increasing sample number in terms of the number of strains resolved, but reassuringly even when only a small number of samples are available we are still able to accurately predict (with 0.068% per base error rate) strains, and when ten or more samples are available we obtain error rates below 0.05% *i*.*e*. 1 error in every 2000 bps from short read data. This is better performance than the state of the art, and sufficiently accurate for high resolution phylogenetics. Strains are resolved more accurately as they increase in coverage (see Figure 4), and in fact, when coverages exceed twenty fold we can resolve strains very reliably, with just 0.011% error rate averaged across strains in the ten sample synthetic data set. We believe therefore that this pipeline will be useful whenever high quality *de novo* strains are required from metagenome short read time series.

This is to our knowledge the first algorithm capable of constructing strains from metagenomes using assembly graphs from multi-sample coassemblies. Graph-based haplotype resolution has been applied to viruses [4] and for eukaryotic transcripts [5, 6], but ours is the first algorithm to resolve strains across multiple gene subgraphs connected through a contig binning procedure. The BayesPaths algorithm is also a substantial algorithmic advance enabling coverage across multiple samples to be incorporated into a rigorous Bayesian procedure that gives uncertainties in both the paths (*i*.*e*. the sequences) and the strain abundances. This algorithm could be utilised outside of the actual STRONG pipeline in other application areas, for example for finding viral haplotypes.

In addition, to the new strain resolution algorithm, BayesPaths, STRONG incorporates a number of useful tools for large-scale variant graph processing, in particular, the tools for extraction of subgraphs that correspond to individual coding genes and the spades-gsimplifier tool for error correction on those graphs. These can be applied to any graph in the GFA format, and could therefore find applicability outside of the context of our pipeline. This also means that in the future we could add alternative choices for the coaseembly step, for instance replacing metaSPAdes with MEGAHIT [25]. Similarly, we plan to add support for alternative binners to CONCOCT.

Currently, we are restricted to core genes that are single-copy and shared across all strains in a MAG. We can in theory use any such genes, so if a particular MAG is of interest the pipeline could be run with a larger set of COGs that are SCGs for that MAG. There would be a cost in terms of increased running time, which will increase with more genes and unitigs in a roughly linear fashion.

The analysis of a time series from an anaerobic digestor illustrates the practicality of our pipeline on a realistically sized data set. We should note though that to resolve strains on these 304 MAGs took nearly 10 days using 64 threads on a standard bioinformatics server (see Table S3. The AD analysis also demonstrates the importance of strain dynamics in a real microbial community with nearly 40% of MAGs exhibiting strain variation, but this variation was relatively stable compared to the MAG dynamics themselves. If strains are functionally redundant to one another we would expect significant neutral fluctuations over time. Therefore this could be evidence for intra-species niche partitioning.

In general, we found a good correspondence between haplotypes inferred from Nanopore reads and the STRONG predictions in the AD data set. For the most abundant MAG, Bin_72, they matched very closely. In addition, the relative abundances of strains were consistent across the two sequencing technologies, despite the use of different DNA extraction protocols, and the different biases inherent in library preparation and sequencing platforms. These technical elements in the data generation process are known to introduce bias at the species level [12], but our findings suggest that intraspecies abundance may generally be robust against such biases, which makes sense in that all the strains of a species will have similar physical properties and genomic traits.

STRONG is an effective strategy to *de novo* resolve subpopulations at high phylogenetic resolution within MAGs, but as discussed in the Introduction, it is important to add the caveat that the haplotype sequences obtained are not equivalent to those from sequencing cultured isolates, where we can identify the resulting genome with a single organism present in the original community. The metagenome strains, in the best case, will correspond to different modal sequences of the taget species, about which substantial unresolved variation may exist. They will correspond to peaks in the probability distribution of all possible sequence configurations, and as such will provide important insights into the naturally occurring variation, but there remains the question of how to identify and quantify the unresolved variation surrounding those peaks. In the worst case, when STRONG is applied to rapidly recombining microbes, such as those found in the oceans [30], the resulting sequences may not even be real in the sense of characterising any true individual. An additional unaddressed question is how to determine when this has occurred, for now we would simply urge caution when using STRONG in cross-sectional studies of rapidly evolving microbes, and suggest that the term ‘metagenome strain’ or ‘metagenome haplotype’ be used when referring to the output sequences. The same caveat does of course apply to any current purely bioinformatics strategy for *de novo* resolution of genomes from metagenomes. Even if a single sample is used for binning and there are no subpopulations, the resulting MAG is still a composite and not a strain in the traditional microbiological sense [39].

An obvious extension of our algorithm would be to resolve the accessory genome into strain genomes. This could be done on a per gene basis by relaxing the requirement that every strain passes through every gene, but an approach that incorporates the path structure in the full metagenomic assembly would be more powerful. Use of the full assembly may be possible in an efficient manner by factorising the variational approximation on a per gene basis and allowing the solutions for one gene to depend on the expectations across their neighbours. Or it may be that more computationally tractable versions of the algorithm can be developed that will scale to larger graphs. In any case the issues discussed above of our inferred ‘strains’ containing unresolved variation will become more pertinent when we extend our algorithm to the full genome, and it will be necessary to consider not just the most likely genome associated with a subpopulation but also its variants.

In the future we also plan to directly incorporate long read information into the strain resolution rather than just using it for validation. It was encouraging therefore to see the correspondence in strain frequencies between the two approaches. We are confident that in the near future, through the combination of long reads with methods similar to those we have introduced in STRONG, that complete metagenome *de novo* strain resolution will become a realistic possibility.

## Conclusion

We have introduced a complete bioinformatics pipeline, STrain Resolution ON assembly Graphs (STRONG), that is capable of extracting single-copy core gene variant graphs from short read metagenome coassemblies for individual metagenome assembled genomes (MAGs). We demonstrated how these graphs and associated per-sample unitig coverages can be used in anovel Bayesian algorithm, BayesPaths, to find MAG strain number, haplotypes and abundances. This approach achieves superior accuracy to variant based methods on synthetic communities and predictions on real data that match those from long Nanopore long reads.

STRONG is freely available from https://github.com/chrisquince/STRONG.

## Methods

### Synthetic data set generation

The *in silico* synthetic communities were generated by first downloading a list of complete bacterial genomes from the NCBI and selecting species with multiple strains present. Genomes were restricted to those that were full genome projects, possessed at least 35 of 36 single-copy core genes (SCGs) identified in [3], and with relatively few contigs (< 5) in the assemblies. Communities were created by specifying species from this list and the number of strains desired. The strains selected were then chosen at random from the candidates for each species, with the extra restrictions that all strains in a species were at least 0.05% and no more than 5% nucleotide divergent on the SCGs from any other strain in the species. This corresponds to a minimum divergence of approximately 15 nucleotides over the roughly 30 kbp region formed by summing the SCGs. The genomes used are given in Tables S1 and S2.

Each species indexed *i* was then given an abundance, *y*_*i,s*_, in each sample, *s* = 1, …, *S*, which was drawn from a lognormal distribution with a species dependent mean and standard deviation, themselves drawn from a normal and gamma distribution respectively:

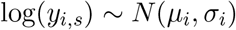

where:

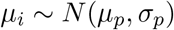

and:

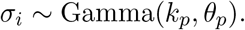

For all four community configurations — *S* equal to 3, 5, 10 and 15 — we used *µ*_*p*_ = 1, *σ*_*p*_ = 0.125, *k*_*p*_ = 1 and *θ*_*p*_ = 1. The species abundances were then normalised to one 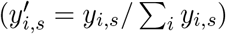. For each strain within a species its proportion in each sample was then drawn from a Dirichlet:

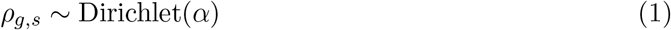

with *α* = 1.

This allowed us to specify a copy number for each genome *g* in species *i* in each sample as 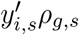. We then generated 150 million paired-end 2×150 bp reads in total across all samples with Illumina HiSeq error distributions using the ART read simulator [21]. The code for the synthetic community generation is available from https://github.com/chrisquince/STRONG_Sim.

### Synthetic data set evaluation

We can determine which contig derived from which reference genome by considering the simulated reads that map onto it. We know which reference each of these came from, enabling us to assign a contig to a genome as that which a majority of its reads derive from. We can then assign each MAG generated by STRONG to a reference species as the one which the majority of its contig’s derive from weighted by the contig length.

### Anaerobic digester sampling and sequencing

#### AD sample collection

We obtained ten samples from a farm anaerobic digestion bioreactor across a period of approximately one year. The sampling times, metadata and accession numbers are given in Table S4. The reactor was fed on a mixture of slurry, whey and crop residues, and operated between 35-40°C, with mechanical stirring. Biomass samples were taken directly from the AD reactor by the facility operators and shipped in ice-cooled containers to the University of Warwick. Upon receipt, they were stored at 4°C and then sampled into several 1-5mL aliquots within a few days. DNA was usually extracted from these aliquots immediately but some were first stored in a −80°C freezer until subsequent thawing and extraction.

#### AD short read sequencing

DNA extraction was performed using the Qiagen Powersoil extraction kit following the manufacturer’s protocol. DNA samples were sequenced by Novogene using the NovaSeq platform with 2×150 bp reads at a mean depth of 11.63 Gbp.

#### AD long read sequencing

Anaerobic digester samples were stored in 1.8 mL Cryovials at −80°C. Samples were defrosted at 4°C overnight prior to DNA extraction. DNA was extracted from a starting mass of 250 mg of anaerobic digester sludge using the MP Biomedical™ FastDNA™ SPIN Kit for Soil (cat no: 116560200) and a modified manufacturers protocol. Defrosted samples were homogenised by pipetting and then transferred to a MP bio™ lysing matrix E tube (cat no: 116914050-CF). Samples were resuspended in 938 *µL* of Sodium phosphate buffer (cat no: 116560205).

Preliminary cell lysis was undertaken using lysozyme at a final concentration of 200 *ng/µL* and 20 *µL* of Molzyme Bug Lysis*™* solution. Samples were mixed by inversion and incubated at 37°C for 30 min on a shaking incubator (< 100 rpm). Lysozyme was inactivated by adding 122 *µL* of MP bio MT buffer and mixing by inversion. Samples were then mechanically lysed in a VelociRuptor V2 bead beating machine (cat no: SLS1401) at 5 *m/s* for 20 seconds then placed on ice for five minutes.

Samples were centrifuged at 14000 g for five minutes to pellet the particulate matter and the supernatant was transferred to a new 1.5 mL microfuge tube. Proteins were precipitated from the crude lysate by adding 250 *µL* of PPS™ (cat no: 116560203) and then mixing by inversion. Precipitated proteins were pelleted for five minutes at 14000 g and the supernatant was transferred to 1000 *µL* of pre mixed DNA binding matrix solution (cat no: 116540408). Samples were mixed by inversion for two minutes.

DNA binding matrix beads were recovered using the MP bio™ spin filter (cat no: 116560210) and manufacturer based spin protocol. The binding matrix was washed of impurities by complete resuspension in 500 *µL* of SEWS-M solution (cat no: 116540405) and centrifuged at 14000 g for five minutes. The DNA binding matrix was then washed for a second time by resuspension in 500 *µL* of 80% EtOH followed by centrifugation at 14000 g for five minutes. Flow though was discarded and centrifuged at 14000 g for two minutes to remove residual EtOH. The binding matrix was left to air dry for 2 minutes then DNA was eluted using 100 *µL* of DES elution buffer at 56°C. Elute was collected by centrifugation at 14000 g for 5 minutes and stored at 4°C prior to library preparation. Eluted DNA concentration was estimated using a Qubit 4™ fluorometer with the dsDNA Broad Range sensitivity assay kit (cat no: Q32853). 260:280 and 260:230 purity ratios were quantified using a Nanodrop™ 2000.

A 1x SPRI clean up procedure was undertaken prior to library construction to further reduce contaminant carry over. Input DNA was standardised to 1.2 *µg* in 48 *µL* of H_2_O using a qubit 4™ fluorometer and dsDNA 1x High Sensitivity assay kit (cat no: Q33231). Library preparation was undertaken using the Oxford Nanopore© Ligation Sequencing Kit (SQK-LSK109) with minor modifications to the manufacturer protocol. The FFPE/End repair incubation step was extended to 30 min at 20°C and 30 min at 65°C, while DNA was eluted from SPRI beads at 37°C for 30 min with gentle agitation. The SQK-LSK109 long fragment buffer was used to ensure removal of non-ligated adaptor units and reduce short fragment carryover into the final sequencing library. The final library DNA concentration was standardised to 250 ng in 12 *µL* of EB using a qubit 4™ fluorometer and dsDNA 1 x High Sensitivity assay kit.

Sequencing was undertaken for 72 hours on an Oxford Nanopore© R 9.4.1 (FLO-MIN106) flow cell with 1489 active pores. DNA was left to tether for 1 hour prior to commencing sequencing. The flow cell and sequencing reaction was controlled by a MinION™ MKII device and the GUI MinKNOW V. 19.12.5. ATP refuelling was undertaken every 18 hours with 75 *µL* of flush buffer (FB). Post Hoc basecalling was undertaken using Guppy V. 3.5.1 and the high accuracy configuration (HAC) mode.

### STRONG pipeline

STRONG processes co-assembly graph regions for multiple metagenomic datasets in order to simultaneously infer the composition of closely related strains for a particular MAG and their core gene sequences. Here, we provide an overview of STRONG. We start from a series of *S* related metagenomic samples, e.g. samples of the same (or highly similar) microbial community taken at different time points or from different locations.

The Snakemake based pipeline begins with the recovery of metagenome-assembled genomes (MAGs) [22]. We perform co-assembly of all available data with the metaSPAdes assembler [28], and then bin the contigs obtained by composition and coverage profiles across all available samples with CONCOCT [3]. Each bin is then analyzed for completeness and contamination based on single-copy core genes, and poor quality bins are discarded. The default criterion is that a MAG requires greater than or equal to 75% of the SCGs in a single copy. While we currently focus on this combination of software tools, in principle we could use any other software or pipeline for MAG recovery, e.g. we could use MEGAHIT as the primary assembler [25] or alternative binning tools or their combination. For each MAG we then extract the full or partial sequences of the core genes that we further refer to as single-copy core gene (SCG) sequences.

The final coassembly graph produced by metaSPAdes cannot be used for strain resolution because, as with other modern assembly pipelines, variants between closely related strains will be removed during the graph simplification process. Instead, we output the initial graph for the final K-mer length used in the (potentially) iterative assembly following processing by a custom executable — spades-gsimplifier based on the SPAdes codebase — to remove likely erroneous edges using a ‘mild’ coverage threshold and a tip clipping procedure. We refer to the resulting graph as a high-resolution assembly graph or HRAG.

The graph edges are then annotated with their corresponding sequence coverage profiles across all available samples. As is typical in de Bruijn graph analysis, the coverage values are given in terms of the k-mer rather than nucleotide coverage. Profile computation is performed by a second tool for aligning reads onto the HRAG: unitig-coverage. The potential advantage of this approach in comparison to estimation based on k-mer multiplicity, is that it can correctly handle the results of any bubble removal procedure that we might want to add to the preliminary simplification phase in future.

For each detected SCG sequence (across all MAGs) we next try to identify the subgraph of the HRAG encoding the complete sequences of all variants of the gene across all strains represented by the MAG. The procedure is described in more detail in the next section. During testing we faced two types of problems here: (1) related strains might end up in different MAGs and (2) some subgraphs might consist of fragments corresponding to several different species. We take several steps to mitigate those problems. Firstly, we compare SCG graphs between all bins, not just MAGs. If an SCG graph shares unitigs between bins, then it is flagged as overlapping. If multiple SCG graphs between MAGs (*>* 10) overlap then we merge those MAGs, combining all graphs and processing them for strains together. Following merging any MAG SCG graphs with overlaps remaining are filtered out and not used in the strain resolution.

After MAG merging and COG subgraph filtering we process the remaining MAGs one by one. Before the core ‘decomposition’ procedure is launched on the set of SCG subgraphs corresponding to a particular MAG, they are subjected to a second round of simplification, parameterised by the mean coverage of the MAG, to filter nodes that are likely to be noise again by the spades-gsimplifier program. This module is described in more detail below. The resulting set of simplified SCGs of the HRAG for a MAG are then passed to the core graph decomposition procedure, which uses the graph structure constraints, along with coverage profiles associated with unitig nodes, to simultaneously predict: the number of strains making up the population represented by the MAG; their coverage depths across the samples; paths corresponding to each strain within every subgraph (each path encodes a sequence of the particular SCG instance).

A fraction of the SCGs in a MAG may properly derive from other organisms due to the possibility of incorrect binning *i*.*e contamination*. In fact, the default 75% single-copy criterion allows up to 25% contamination. In addition, the subgraph extraction is not always perfect. We therefore add an extra level of filtering to the BayesPaths algorithm, iteratively running the program for all SCGs, but then filtering those with mean error rates, defined by Equation 18, that exceed a default of 2.5 times the median deviation from the median gene error. Filtering on the median deviation is in general a robust strategy for identifying outliers. As a result of this filtering the pipeline only infers strain sequences on a subset of the input SCGs. We have found, however, that the number of SCGs for which strain haplotypes are inferred is sufficient for phylogenetics.

### Relevant subgraph extraction

Provided with the predicted (partial) gene sequence, 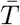, and the upper bound on the length of the coding sequence, *L*, defined as 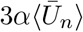 where 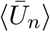 is the average length in amino acids of that SCG in the COG database, and *α* = 1.5 by default. The procedure for relevant HRAG subgraph extraction involves the following steps. First, the sequence 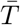 is split into two halves, 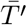 and 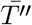, keeping the correct frame (both 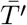 and 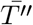 are forced to divide by 3).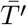 and 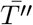 are then processed independently. Without loss of generality we describe the processing of 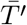

1. Identify the path 𝒫 corresponding to 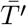 in the HRAG. We denote its length as *L*_*P*_.
2. Launch a graph search of the stop codons to the right (left) of the rightmost (leftmost) position of 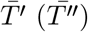. The stop codon search is frame aware and is performed by a depth-first search (DFS) on the graph in which each vertex corresponds to a pair of the HRAG position and the partial sequence of the last traversed codon ^1^. Vertices of this ‘state graph’ are naturally connected following the HRAG constraints. The search is cut off whenever a vertex with a frame state encoding a stop codon sequence is identified. Several stop codons can be identified within the same HRAG edge sequence in ‘different frames’, moreover the procedure correctly identifies all stop codons even if the graph contains cycles (although such subgraphs may be ignored in later stages of the pipeline).
3. The ‘backward’ search of the stop codons ‘to the left’ is actually implemented as a ‘forward’ search of the complementary sequences from the complementary position in the graph. Note that, as in classic ORF analysis, while the identified positions of the stop codons ‘to the right’ correspond to putative ends of the coding sequences for some of the variants of the analyzed gene, positions of the stop codons ‘to the left’ only provide the likely boundary for where the coding sequence can start. In particular, left stop codons are likely to fall within the coding sequence of the neighbouring gene (in a different frame). Actual start codons are thus likely to lie somewhere on the path (with sequence length divisible by 3) between one of the ‘left’ stop codons and one of the ‘right’ stop codons. For reasons of simplicity, further analysis of edges on the paths between left (right) stop codons ignores the constraint of divisibility by 3.
4. After the sets of ‘left’ and ‘right’ stop codon positions are identified along with the shortest distances between them and the 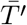 path, we attempt to gather the relevant subgraph given by the union of edges lying on some path of a constrained length (see further) between some pair of left and right stop codons. First, for each pair (*s, t*) of the left and right stop codon positions we compute the maximal length of the paths that we want to consider *L*_*s,t*_ as *L*_*P*_ + min dist from s to start of 𝒫 + min dist from end of 𝒫 to t. The edge *e* = (*v, w*) is considered relevant if there exists a pair of left (right) stop codon positions (*s*′, *t*′) such that the edge *e* lies on the path of length not exceeding *L*_*s,t*_ between *s*′ and *t*′, which is equivalent to checking that min dist(*s*′, *v*) + length(*e*) + min dist(*w, t*′) < *L*_*s,t*_. To allow for efficient checks of the shortest distances we precompute them by launching the Dijkstra algorithm from all left (right) stop codon positions in the forward (backward) direction ^2^.
5. We then exclude from the set of relevant edges the edges that are too far from any putative (right) stop codon to be a part of any COG instance. In particular, we exclude any edge *e* = (*v, w*), such that the minimal distance from vertex *w* to any of the right stop codon positions exceeds *L*.
6. After the sets of the graph edges potentially encoding the gene sequence are gathered for 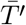 and 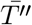 the union of the two sets, *εℛ*, is then taken and augmented by the edges, connecting the ‘inner’ *εℛ* vertices (vertices which have at least one outgoing and at least one incoming edge in the *εℛ*) to the rest of the graph.

Initial splitting of 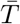 into 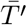 and 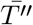 is required to detect relevant stop codons which are not reachable from the last position of 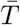 in HRAG (or from which the first position of 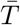 in HRAG can not be reached). In addition to the resulting component in gfa format, we also store the positions of the putative stop codons, and ids of edges connecting the component to the rest of the graph (further referred to as ‘sinks’ and ‘sources’).

### Subgraph simplification

While processing SCG subgraphs from a particular MAG we use the available information on the coverage of the MAG in the dataset. In particular, we set up the simplification module to remove tips (a node with no successors) below a certain length and edges with coverage below a fraction of the total coverage across all samples. If a tip is not removed it is labelled as a ‘dead-end’ to distinguish it from potential connections to the rest of the graph.

While simplifying a COG subgraph, edges connecting it to the rest of the assembly graph should be handled with care (in particular, they should be excluded from the set of potential tips). This is because in the BayesPaths algorithm they form potential sources and sinks of the possible haplotype paths. Moreover, during the simplification the graph changes, and such edges might become part of longer edges. Since we are interested in which dead-ends of the component do, and do not lead to the rest of the graph, the output contains the up-to-date set of connections of the simplified component to the rest of the graph.

We now briefly describe the implemented procedures based on ‘relative coverage’ criteria. Amongst other procedures for erroneous edge removal SPAdes implements a procedure considering the ratio of the edge coverage to the adjoining coverage of edges adjacent to it. We define an edge *e* as ‘predominated’ by vertex *v* incident to it if there is edge *e*_1_ outgoing from *v* and edge *e*_2_ incoming to *v* whose coverages exceed the coverage of *e* at least by a factor of *α* (by default equal to 5). Short edges (shorter than *k* + *ϵ*) predominated by both vertices incident to them are then removed from the graph. Erroneous graph elements in high genomic coverage graph regions often form subgraphs of three or more erroneous edges. SPAdes implements a procedure for search (and subsequent removal) of subgraphs limited by a set of predominated edges. Starting from a particular edge (*v, w*) predominated by vertex *v*, the graph is traversed from vertex *w* breadth-first without taking into account the edge directions. If the vertex considered at the moment predominates the edge by which it was entered, the edges incident to it are not added to the traversal. The standard limitation of erroneous edge lengths naturally transforms into a condition of maximum length of the path between the vertices of the traversed subgraph. A limit on the maximum total length of its edges is additionally introduced.

### BayesPaths

#### The model

We define an assembly graph *𝒢* = (*𝒱, ε*) as a collection of unitig sequence vertices *𝒱* = 1, …, *V* and directed edges *ε* ⊆ *𝒱* × *𝒱*. Each edge defines an overlap and comprises a pair of vertices and directions (*u*^*d*^ → *v*^*d*^) ∈ *ε* where *d ∈* {+, −} and indicates whether the overlap occurs between the sequence (+) or its reverse complement (−). We define

- Counts *x*_*v,s*_ for each unitig *v* = 1, …, *V* in sample *s* = 1, …, *S*
- Paths for strain *g* = 1, …, *G* defined by 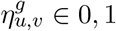 indicating whether strain *g* passes through that edge in the graph
- Flow of strain *g* through unitig 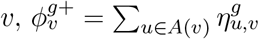 and 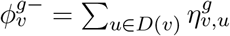 where *A*(*v*) is the set of ancestors of *v* and *D*(*v*) descendants in the assembly graph
- The following is true 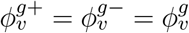
- Strain intensities *γ*_*g,s*_ as the rate per position that a read is generated from *g* in sample *s*
- Unitig lengths *L*_*v*_
- Unitig bias *θ*_*v*_ is the fractional increase in reads generated from *v* given factors such as GC content influencing coverage
- Source node *s* and sink node *t* such that 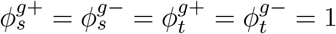

Then assume normally distributed counts for each node in each sample giving a joint density for observations and latent variables:

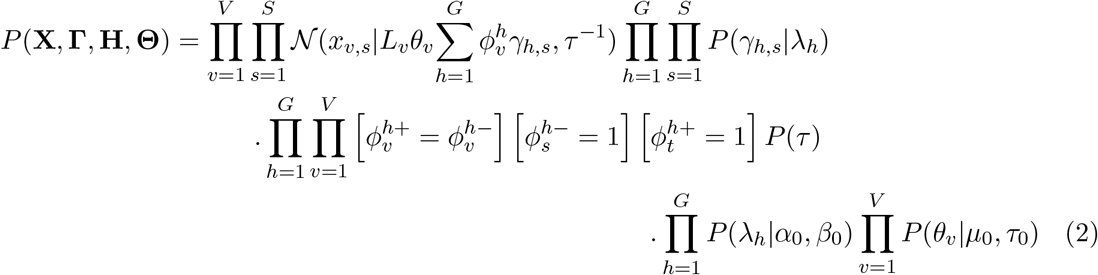

where [] indicates the Iverson bracket evaluating to 1 if the condition is true and zero otherwise. We assume an exponential prior for the *γ*_*g,s*_ with a rate parameter that is strain dependent, such that:

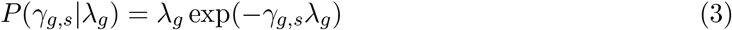

We then place gamma hyper-priors on the *λ*_*g*_:

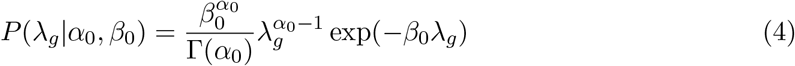

This acts as a form of automatic relevance determination (ARD) forcing strains with low intensity across all samples to zero in every sample [8].

We use a standard Gamma prior for the precision:

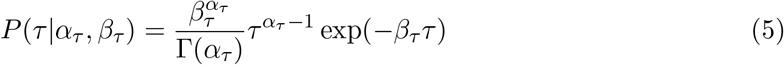

For the biases *θ*_*v*_ we use a truncated normal prior:

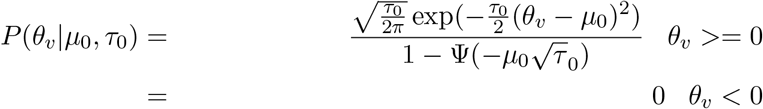

where Ψ is the standard normal cumulative distribution. The mean of this is set to one, *µ*_0_ = 1, so that our prior is that the coverage on any given node is unbiased, with a fairly high precision *τ*_0_ = 100, to reflect an assumption that the observed coverage should reflect the summation of the strains. Finally, we assume a uniform prior over the possible discrete values of the 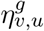 If the assembly graph is a directed acyclic graph (DAG) then 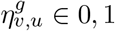. We have found that for most genes and typical kmer lengths this is true, but we do not need to assume it.

### Variational Approximation

We use variational inference to obtain an approximate solution to the posterior distribution of this model [7]. Variational inference is an alternative strategy to Markov chain Monte Carlo (MCMC) sampling. Rather than attempting to sample from the posterior distribution, variational inference assumes a tractable approximating distribution for the posterior, and then finds the parameters for that distribution that minimise the Kullback-Leibler divergence between the approximation and the true posterior distribution. Further, in mean-field variational inference the approximation can be factorised into a product over a number of components that each approximate the posterior of a parameter in the true distribution. In practice the Kullback-Leibler divergence is not computable because it depends on the evidence, i.e. the joint distribution marginalised over all latent variables. Instead, inference is carried out by maximising the evidence lower bound (ELBO), which is equal to the negative of the Kullback-Leibler divergence plus a constant, that constant being the evidence. In our case, because all the distributions are conjugate we can employ CAVI, coordinate ascent variational inference, to iteratively maximise the ELBO.

Our starting point is to assume the following factorisation for the variational approximation:

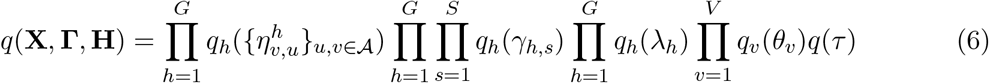

where 𝒜 is the set of edges in the assembly graph and 𝒱 = 1, …, *V* the set of unitig sequence vertices. Note that we have assumed a fully factorised approximation except for the 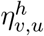 the paths for each strain through the graph. There we assume that the path for each strain forms a separate factor allowing strong correlations between the different elements of the path. This is therefore a form of structured variational inference [20].

To obtain the CAVI updates we use the standard procedure of finding the log of the optimal distributions *q* for each set of factorised variables as the expectation of the log joint distribution Equation 2 over all the other variables, except the one being optimised. Using an informal notation we will denote these expectations as ⟨ln *P* ⟩_*− q*_*j* where *q*_*j*_ is the variable being optimised.

Then the mean field update for each set of 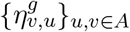 is derived as:

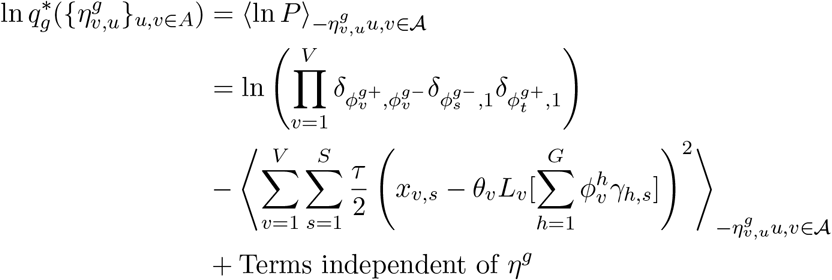

Consider the second term only:

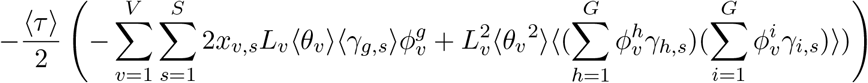

This becomes:

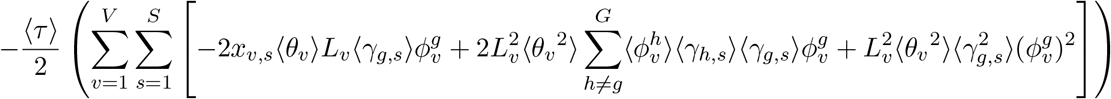

Which can be reorganised to:

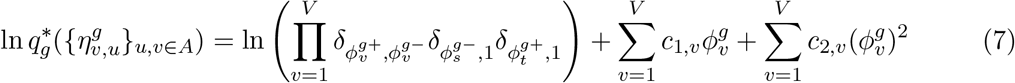

Where:

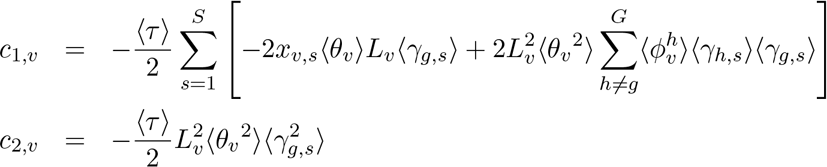

It is apparent from Equation 7 that the 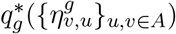 takes the form of a multivariate discrete distribution with |*u, v ∈ A*| dimensions. The first term in Equation 7 enforces the flow constraints, and does not separate across nodes, the next two terms are effectively coefficients on the total flow through a unitig and its square. The updates for the other variables below, depend on the expected values of the total flow through each of the unitig nodes for the strain *g*, 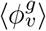, which themselves depend on the 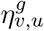 These expected values can be efficiently obtained for all *v* by representing Equation 7 as a factor graph comprising nodes consisting of factors corresponding to both the constraints and the flow probabilities through each node with variables 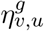. We can then find the marginal probabilities for both the 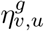 and the 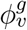 using the Junction Tree algorithm [41], from these we can calculate the required expectations. Next we consider the mean field update for the *γ*_*g,s*_:

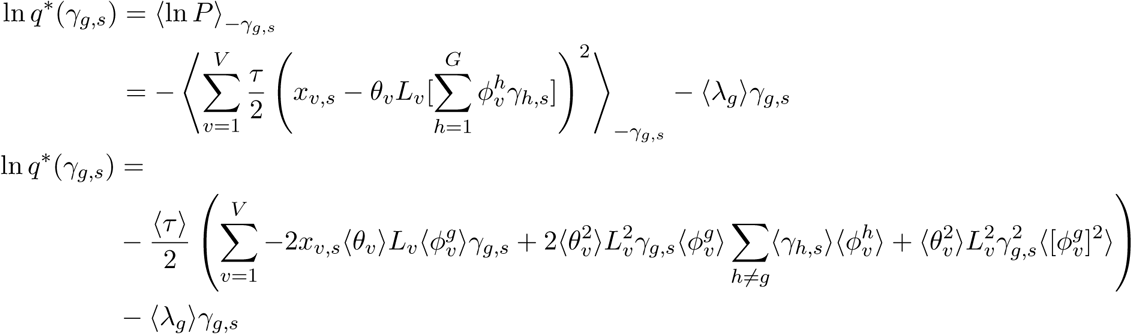

with the restriction *γ*_*g,s*_ *>* 0 this gives a normal distribution but truncated to (0, inf) for *γ*_*g,s*_, with mean and precision:

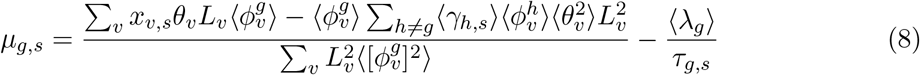

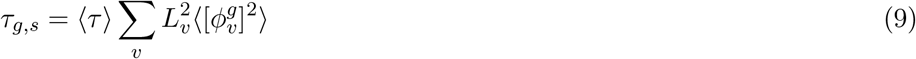

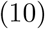

Derivations for the other updates follow similarly giving a Gamma posterior for the *τ* with parameter updates:

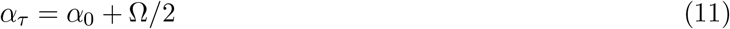

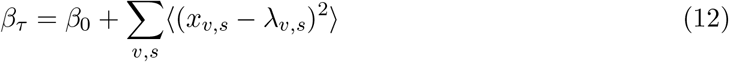

where Ω = *VS* and we have used *λ*_*v,s*_ as a short hand for the predicted count number:

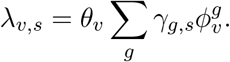

Then the *τ* have the following expectations and log expectations:

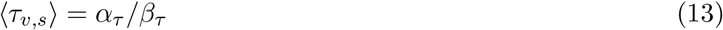

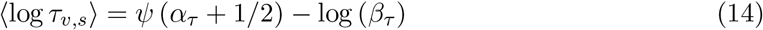

where *Ψ* is the digamma function. The biases *θ*_*v*_ have a truncated normal distribution and their updates can be derived similar to the above.

### Evidence lower bound (ELBO)

Iterating the CAVI updates defined above will generate a variational approximation that is optimal in the sense of maximising the evidence lower bound (ELBO) so called because it bounds the log evidence, *log*(*p*(*x*)) ≥ *ELBO*(*q*(*z*)). It is useful to calculate the ELBO whist performing CAVI updates to verify convergence and the ELBO itself is sometimes used as a Bayesian measure of model fit, although as a bound that may be controversial [7]. The ELBO can be calculated from the relationship:

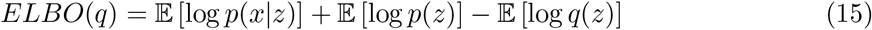

The first term is simply the expected log-likelihood of the data given the latent variables. In our case it is:

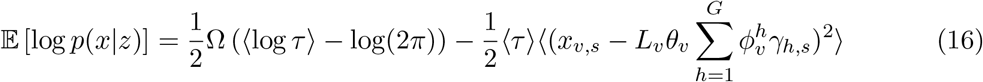

where Ω = *V S* and the expectations are over the optimised distributions *q*.

The second term is the expectation under *q*(*z*) of the log prior distributions. In our case with standard distributions it is easy to calculate for instance for each of the *γ*_*g,s*_:

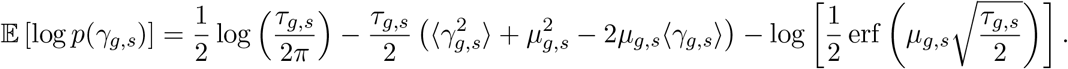

With the *µ*_*g,s*_ and *τ*_*g,s*_ given by their current values derived from Equation 10 and the moments of *γ*_*g,s*_ calculated from a truncated normal distribution with those current parameters. The third terms are simply the negative entropy of the variational approximations and for the standard distributions used here are easily calculated.

### Implementation details

One update of the algorithm consists of updating each variable or sets of variables in turn given the current optimal solutions of the other distributions. In practice we update:

- Compute the marginal flows 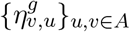 for each strain *g* = 1, …, *G* in turn using Equation 7 and the Junction Tree algorithm. This can be performed for each single copy-core gene independently
- Update the truncated normal strain abundances *q*(*γ*_*g,s*_) for each strain in each sample, *s* = 1, …, *S* using Equation 10
- Update the *q*(*τ*)
- Update the ARD parameter distributions *q*(*λ*_*g*_) if used
- Update the nodes biases *q*(*θ*_*v*_)
- Check for redundant or low abundance strains and remove (see below)

After a maximum fixed number of iterations or if the ELBO converges we stop iterating. Variational inference can be sensitive to initial conditions as it can only find local maxima of the ELBO, we therefore use a previously published variational Bayesian version of non-negative matrix factorisation [8], to find an initial approximate solution.

### Empirical modelling of node variances

For low-coverage MAGs a precision that is identical for all nodes performs satisfactorily, but since the true distribution of read counts is Poisson this overestimates the precision for nodes with high counts *x*_*v,s*_. To address we developed an empirical procedure where we first calculate ⟨log *τ*_*v,s*_) for each node using Equation 14 as:

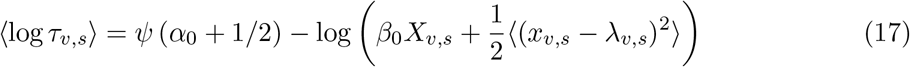

a quantity which exhibits high variability, so we then smooth this over log(*x*_*v,s*_) using generalised additive models as implemented in pyGAM [36] to give ⟨log *τ*_*v,s*_⟩*∗*. The term *β*_0_*X*_*v,s*_ gives a prior which is effectively Poisson. We then obtain ⟨*τ*_*v,s*_⟩ as exp(⟨log *τ*_*v,s*_⟩*∗*). This procedure has no theoretical justification but gives good results in practice. This approach of modelling a non-Gaussian distribution as a Gaussian with empirical variances is similar to that used in voom for RNASeq [23].

### Cross-validation to determine optimum number of strains

The ARD procedure usually converges on the correct number of strains except for high-coverage MAGs where overfitting may occur and too many strains can be predicted. We therefore additionally implemented a cross-validation procedure, splitting the entire data matrix *x*_*v,s*_ into test and train folds (default ten folds) and training the model on the train fold and then testing on the held out data. The correct number of strains was then taken to be the one that maximised the log predictive posterior with an empirical variance reflecting the Poisson nature of the data. The exact test statistic being:

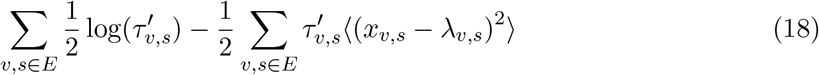

Where 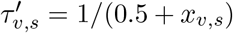 and *E* indicates data points in the test set to down-weight high read count nodes reflecting approximately Poisson noise.

### Nanopore sequence analysis

#### Sequence preprocessing

To enable a qualitative comparison between haplotypes obtained from the Nanopore reads and the BayesPaths predictions we developed the following pipeline applied at the level of individual single-copy core genes (SCGs) from MAGs:

1. We mapped all reads using minimap2 [26] against the SCG contig consensus ORF sequence and selected those reads with alignments that spanned at least one third of the gene with a nucleotide identity *>* 80%.
2. We then extracted the aligned portion of each read, reverse complementing if necessary, so that all reads ran in the same direction as the SCG ORF.
3. We then obtained the variant positions on the consensus from the output of the DESMAN pipeline [32]. These are variant positions prior to haplotype calling representing the total unlinked variation observed in the short reads.
4. For each Nanopore fragment we aligned against the SCG ORF using a Needleman-Wunsch global alignment and generated a condensed read comprising bases only from the short read variant positions.

This provided us with a reduced representation of each Nanopore read effectively filtering variation that was not observed in the short reads. These reduced representations were then used to calculate distances, defined as Hamming distances on the variant positions normalised by number of positions observed, both between the reads and between the reads and the predicted COG sequences from BayesPaths. From these we generated NMDS plots indicating sequence diversity, and they provided an input to the hybrid Nanopore strain resolution algorithm below.

#### EM algorithm for hybrid Nanopore strain resolution

We also developed a simple EM algorithm for direct prediction of paths and their abundances on the short read assembly graph that are consistent with a set of observed long reads. We began by mapping the set of *n* = 1, …, *N* Nanopore sequences denoted *{*𝒮_*n*_*}* onto the corresponding simplified SCG graph generated by STRONG using GraphAligner [33]. This provided us with *N* optimal alignment paths as walks in our SCG graph. We denote this graph 𝒢 comprising unitig vertices *v* and edges *e* ∈ {*u, v*} defining overlaps.

We assume, as is almost always the case that the graphs contain no unitigs in both forward and reverse configurations, and that there are no cycles, so that each SCG is a directed acyclic graph (DAG) with one copy of each unitig, and we only need to track the direction that each overlap enters and leaves each unitig. Then best alignment walks comprise a sequence of edges, 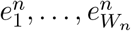 where *W*_*n*_ is the number of edges in the walk of read *n*, that traverse the graph.

Given these observed Nanopore reads we aim to reconstruct *G* haplotypes comprising paths from a dummy source node *s*, which has outgoing edges to all the true source nodes in the graph, through the graph to a dummy sink *t*, which connects all the true sinks. We further assume that each haplotype has relative frequency *π*_*g*_. Each such haplotype path 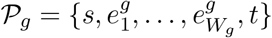 will translate into a nucleotide sequence 𝒯_*g*_. We assume that these haplotypes generate Nanopore reads with a fixed error rate *c* which gives a likelihood:

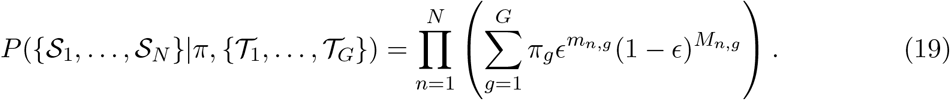

where *m*_*n,g*_ is the number of basepair mismatches between 𝒮_*n*_ and 𝒯_*g*_ counting insertions, deletions and substitutions equally and *M*_*n,g*_ the number of matches.

To maximise this likelihood we used an Expectation-Maximisation algorithm. Iterating the following steps until convergence:

1. E-step: Calculate the responsibility of each haplotype for each sequence as:

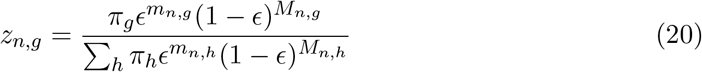

Alignments of reads against haplotypes were performed using vsearch [34].
2. M-step: We update each haplotype by finding the most likely path on the short read graph given the current expectations. These are calculated by assigning a weight 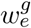 to each edge *e* in the graph as:

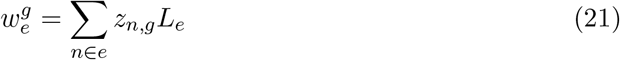

where *n ∈ e* are the set of reads whose optimal alignment contains that edge and *L*_*e*_ is the unique length of the unitig the edge connects to, *i*.*e*. ignoring the overlap length. We then find for haplotype *g* the maximal weight path through this DAG using a topological sort. The error rates are updated as:

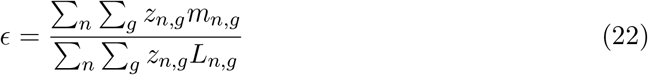

where *L*_*n,g*_ are the alignment lengths.

As is often the case with EM algorithms convergence depends strongly on initial conditions. Therefore we initialise using a partitioning around medoids clustering using the distances calculated in Methods - Nanopore Sequence Analysis. We can estimate the number of haplotypes from the negative log-likelihood as a function of haplotype number.

## Declarations

### Ethics approval and consent to participate

Not applicable.

### Consent for publication

Not applicable.

### Availability of data and materials

The anaerobic digester time series have been uploaded to the ENA as part of the study PRJEB39861. The STRONG pipeline and synthetic community data are available from https://github.com/chrisquince/STRONG and the BayesPaths algorithm https://github.com/chrisquince/BayesPaths. The code for synthetic community generation from https://github.com/chrisquince/STRONG_Sim and the Nanopore EM algorithm https://github.com/chrisquince/NanoHap.

### Competing interests

The authors declare that they have no competing interests.

## Funding

This work for made possible through the MRC Methodology Grant ‘Strain resolved metagenomics for medical microbiology’ MR/S037195/1. CQ is also funded through MRC fellowship (MR/M50161X/1) as part of the CLoud Infrastructure for Microbial Genomics (CLIMB) consortium (MR/L015080/1). SR is funded through BBSRC ‘EBI Metagenomics - enabling the reconstruction of microbial populations’ (BB/R015171/1). OSS acknowledges funding through the BBSRC (BB/N023285/1 and BB/L502029/1).

## Authors’ contributions

CQ devised and coded the BayesPaths algorithm, assisted with the creation of the STRONG pipeline, analysed results and wrote the MS. SN devised and coded the graph algorithms, assisted with the creation of the STRONG pipeline, and edited the MS. SR coded the STRONG pipeline. RJ generated and processed the AD Nanopore sequence data. OSS helped devise the AD sequencing study. JKS assisted with the STRONG pipeline and edited the MS. AL contributed to the creation of the algorithms and edited the MS. AME contributed to the creation of the algorithms and assisted with figures. RC tested the STRONG pipeline, contributed to the creation of the algorithms and edited the MS. AED helped plan the STRONG pipeline, assisted the creation of the algorithms and edited the MS.

## Acknowledgements

The subgraph extraction procedure was developed following discussions with Tatiana Dvorkina (PhD student in SPbSU).

## Footnotes

1 Due to the properties of the procedure and the fact that it deals with DBGs, the actual implementation encodes frame state as an integer [0,2] rather than the string of last partially traversed codon.

2 Actually Dijkstra runs are initiated from the ends/starts of corresponding edges and the distances are later corrected.

## Notes

https://github.com/chrisquince/STRONG

https://github.com/chrisquince/BayesPaths

### Competing Interest Statement

The authors have declared no competing interest.

