## Supplementary Tables and Figures for "Metagenomics Strain Resolution on Assembly Graphs"

### Supporting information

#### Supplementary tables

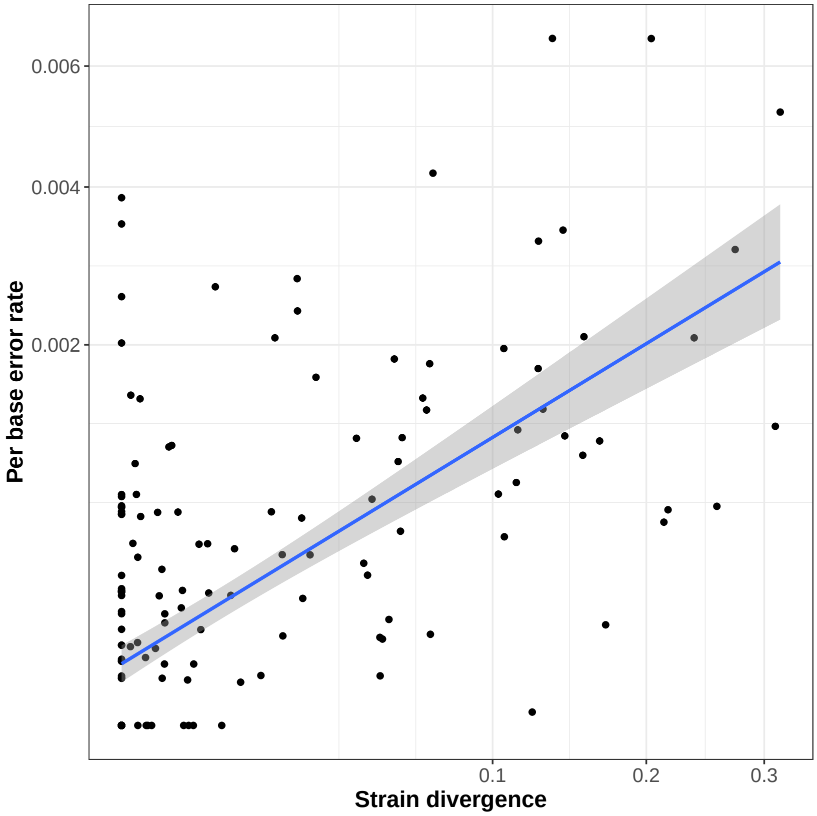


Figure S1: **Correlation between strain divergence and error rate for the synthetic community data.** For the four synthetic community data sets combined, Synth_03, Synth_05, Synth_10 and Synth_15, we show the estimated error as path divergence (see main text) against actual nucleotide error rates. The straight line is a linear regression. The error rates and divergences were correlated (Pearson's correlation: '$r = 0.56$, $p < 2.2e - 16$).

| Species name | Tax. ID. | Strain ID | No. contigs | Genbank Accn no. |
|---|---|---|---|---|
| Pseudomonas chlororaphis | 587753 | 1037915 | 1 | GCF˙000264555.1 |
| Pseudomonas chlororaphis | 587753 | 86192 | 1 | GCF˙000761195.1 |
| Pseudomonas chlororaphis | 587753 | 587753 | 1 | GCF˙000698865.1 |
| Pseudomonas chlororaphis | 587753 | 1038921 | 1 | GCF˙000281915.1 |
| Bordetella holmesii | 35814 | 1247648 | 4 | GCF˙000598125.1 |
| Bordetella holmesii | 35814 | 1172205 | 1 | GCF˙000765395.1 |
| Bordetella holmesii | 35814 | 1247646 | 4 | GCF˙000572015.1 |
| Bordetella holmesii | 35814 | 1266729 | 2 | GCF˙000341465.1 |
| Comamonas testosteroni | 285 | 399795 | 1 | GCF˙000168855.1 |
| Comamonas testosteroni | 285 | 1392005 | 1 | GCF˙000739375.1 |
| Acetobacter pasteurianus | 438 | 438 | 1 | GCF˙001183745.1 |
| Rhodococcus erythropolis | 1833 | 234621 | 4 | GCF˙000010105.1 |
| Rhodococcus erythropolis | 1833 | 1136179 | 2 | GCF˙000454045.1 |
| Rhodococcus erythropolis | 1833 | 1289591 | 3 | GCF˙000696675.1 |
| Rickettsia prowazekii | 782 | 272947 | 1 | GCF˙000195735.1 |
| Clostridium sporogenes | 1509 | 1509 | 1 | GCF˙001020205.1 |
| Clostridium sporogenes | 1509 | 471871 | 2 | GCF˙000155085.1 |
| Mycoplasma capricolum | 2095 | 1124992 | 1 | GCF˙000835085.1 |
| Mycoplasma capricolum | 2095 | 40480 | 1 | GCF˙000953375.1 |
| Achromobacter xylosoxidans | 85698 | 1167634 | 1 | GCF˙000967095.2 |
| Achromobacter xylosoxidans | 85698 | 562971 | 1 | GCF˙000758265.1 |
| Sinorhizobium meliloti | 382 | 1230587 | 4 | GCF˙000304415.1 |
| Sinorhizobium meliloti | 382 | 698936 | 3 | GCF˙000147775.2 |
| Mycoplasma gallisepticum | 2096 | 1159201 | 1 | GCF˙000286755.1 |
| Mycoplasma gallisepticum | 2096 | 710128 | 1 | GCF˙000025365.1 |
| Mycoplasma gallisepticum | 2096 | 708616 | 1 | GCF˙000025385.1 |
| Riemerella anatipestifer | 34085 | 1228997 | 1 | GCF˙000295655.1 |
| Riemerella anatipestifer | 34085 | 1271752 | 1 | GCF˙000331695.1 |
| Riemerella anatipestifer | 34085 | 693978 | 1 | GCF˙000183155.1 |
| Riemerella anatipestifer | 34085 | 1455062 | 1 | GCF˙001077795.1 |
| Riemerella anatipestifer | 34085 | 992406 | 1 | GCF˙000191565.1 |
| Leuconostoc mesenteroides | 1245 | 33966 | 2 | GCF˙001047695.1 |
| Bacillus atrophaeus | 1452 | 720555 | 1 | GCF˙000165925.1 |
| Bacillus atrophaeus | 1452 | 1239783 | 1 | GCF˙000385965.2 |
| Lactobacillus fermentum | 1613 | 1381124 | 2 | GCF˙000466785.3 |
| Anaplasma phagocytophilum | 948 | 1184254 | 1 | GCF˙000439795.1 |
| Borreliella garinii | 29519 | 1081646 | 3 | GCF˙000239475.1 |
| Borreliella garinii | 29519 | 1421551 | 1 | GCF˙000691545.1 |
| Borreliella garinii | 29519 | 1234596 | 1 | GCF˙000300045.1 |
| Flavobacterium psychrophilum | 96345 | 96345 | 1 | GCF˙000739395.1 |
| Flavobacterium psychrophilum | 96345 | 96345 | 1 | GCF˙000971645.1 |
| Flavobacterium psychrophilum | 96345 | 1452724 | 1 | GCF˙000754405.1 |
| Coxiella burnetii | 777 | 227377 | 2 | GCF˙000007765.1 |
| Coxiella burnetii | 777 | 434924 | 2 | GCF˙000019885.1 |
| Coxiella burnetii | 777 | 360115 | 2 | GCF˙000018745.1 |
| Bradyrhizobium japonicum | 375 | 375 | 1 | GCF˙000807315.1 |
| Salinispora tropica | 168695 | 369723 | 1 | GCF˙000016425.1 |
| Brucella ovis | 236 | 444178 | 2 | GCF˙000016845.1 |
| Agrobacterium tumefaciens | 358 | 1435057 | 4 | GCF˙000576515.1 |
| Neorhizobium galegae | 399 | 1028801 | 3 | GCF˙000731295.1 |

Table S1: **Genomes used in the synthetic communities (part I).**

| Species name | Tax. ID. | Strain ID | No. contigs | Genbank Accn no. |
|---|---|---|---|---|
| Lactobacillus jensenii | 109790 | 575606 | 4 | GCF˙000161895.2 |
| Corynebacterium glutamicum | 1718 | 1079988 | 2 | GCF˙000233355.2 |
| Haloferax mediterranei | 2252 | 523841 | 4 | GCF˙000306765.2 |
| Haloferax mediterranei | 2252 | 523841 | 4 | GCF˙000685635.1 |
| Haemophilus parainfluenzae | 729 | 862965 | 1 | GCF˙000210895.1 |
| Streptococcus dysgalactiae | 1334 | 617121 | 1 | GCF˙000307185.1 |
| Lactobacillus salivarius | 1624 | 712961 | 4 | GCF˙000143435.1 |
| Yersinia frederiksenii | 29484 | 1454377 | 2 | GCF˙000834215.1 |
| Edwardsiella tarda | 636 | 498217 | 2 | GCF˙000020865.1 |
| Bacillus pumilus | 1408 | 315750 | 1 | GCF˙000017885.1 |
| Bacillus licheniformis | 1402 | 1402 | 1 | GCF˙000876525.1 |
| Bacillus licheniformis | 1402 | 1402 | 4 | GCF˙000948275.1 |
| Saccharolobus solfataricus | 2287 | 2287 | 1 | GCF˙000968435.1 |
| Morganella morganii | 582 | 1124991 | 1 | GCF˙000286435.2 |
| Ralstonia solanacearum | 305 | 859656 | 1 | GCF˙000197855.1 |
| Ralstonia solanacearum | 305 | 1031711 | 2 | GCF˙000215325.1 |
| Ralstonia solanacearum | 305 | 305 | 1 | GCF˙001373335.1 |
| Ralstonia solanacearum | 305 | 305 | 2 | GCF˙001299555.1 |
| Ralstonia solanacearum | 305 | 305 | 1 | GCF˙001373255.1 |
| Pantoea ananatis | 553 | 706191 | 1 | GCF˙000025405.2 |
| Pantoea ananatis | 553 | 932677 | 2 | GCF˙000270125.1 |
| Pantoea ananatis | 553 | 1123863 | 2 | GCF˙000283875.1 |
| Pantoea ananatis | 553 | 1095774 | 2 | GCF˙000233595.1 |
| Bartonella bacilliformis | 774 | 1293904 | 4 | GCF˙000709855.1 |
| Bartonella bacilliformis | 774 | 1293906 | 3 | GCF˙000709775.1 |
| Bartonella bacilliformis | 774 | 1293907 | 4 | GCF˙000709875.1 |
| Bartonella bacilliformis | 774 | 1293910 | 4 | GCF˙000709755.1 |
| Actinobacillus pleuropneumoniae | 715 | 537457 | 4 | GCF˙000020405.1 |
| Actinobacillus pleuropneumoniae | 715 | 416269 | 1 | GCF˙000015885.1 |
| Sulfolobus islandicus | 43080 | 429572 | 1 | GCF˙000022385.1 |
| Sulfolobus islandicus | 43080 | 1132501 | 1 | GCF˙000245175.1 |
| Sulfolobus islandicus | 43080 | 427317 | 1 | GCF˙000022405.1 |
| Sulfolobus islandicus | 43080 | 930943 | 1 | GCF˙000189575.1 |
| Streptococcus thermophilus | 1308 | 1308 | 1 | GCF˙000971665.1 |
| Streptococcus thermophilus | 1308 | 264199 | 1 | GCF˙000011825.1 |
| Streptococcus thermophilus | 1308 | 1187956 | 1 | GCF˙000262675.1 |
| Streptococcus thermophilus | 1308 | 1308 | 1 | GCF˙001008015.1 |
| Streptococcus thermophilus | 1308 | 767463 | 1 | GCF˙000182875.1 |
| Methanosarcina barkeri | 2208 | 1434109 | 2 | GCF˙000969985.1 |
| Methanosarcina barkeri | 2208 | 796385 | 1 | GCF˙001027005.1 |
| Methanosarcina mazei | 2209 | 1434114 | 1 | GCF˙000970225.1 |
| Methanosarcina mazei | 2209 | 1434115 | 1 | GCF˙000970185.1 |
| Methanosarcina mazei | 2209 | 213585 | 1 | GCF˙000970205.1 |
| Methanosarcina mazei | 2209 | 1434117 | 1 | GCF˙000970165.1 |
| Methanosarcina mazei | 2209 | 192952 | 1 | GCF˙000007065.1 |
| Bifidobacterium bifidum | 1681 | 702459 | 1 | GCF˙000165905.1 |
| Bifidobacterium bifidum | 1681 | 883062 | 1 | GCF˙000164965.1 |
| Bifidobacterium bifidum | 1681 | 1681 | 2 | GCF˙001020375.1 |
| Bifidobacterium bifidum | 1681 | 500634 | 1 | GCF˙001025135.1 |
| Bifidobacterium bifidum | 1681 | 398513 | 1 | GCF˙000273525.1 |

Table S2: **Genomes used in the synthetic communities (part II).**

| Data set | Synth_03 | Synth_05 | Synth_10 | Synth_15 | AD |
|---|---|---|---|---|---|
| Size (Gbp) | 45 | 45 | 45 | 45 | 116.3 |
| No. of samples | 3 | 5 | 10 | 15 | 10 |
| No. of MAGs | 34 | 37 | 39 | 40 | 304 |
| Assembly | 16.92 | 15.63 | 14.50 | 9.98 | 49.93 |
| Binning | 0.03 | 0.90 | 0.60 | 0.03 | 77.30 |
| Subgraphs | 0.60 | 1.58 | 0.90 | 0.93 | 35.05 |
| BayesPaths | 27.43 | 35.1 | 36.82 | 19.4 | 228.65 |

Table S3: **Approximate STRONG run times in hours for data sets in this study.** We give time in hours for the major steps of the STRONG pipeline. These were run using 64 cores of a 192 core Linux x86_64 server running Intel(R) Xeon(R) CPUs E7-8850 v2 @ 2.30GHz.

| Sample | Week | Nucleotides | Temperature | H2S | CH4 | O2 |
|---|---|---|---|---|---|---|
| AD7˙W5˙Repeat2 | 5 | 1.48e+10 | 38.6 | 71.6 | 55.6 | 0.616 |
| AD7˙W10 | 10 | 1.34e+10 | 36.8 | 267 | 52.9 | 0.299 |
| AD7˙W14 | 14 | 1.15e+10 | 39 | 111 | 63.3 | 0.499 |
| AD7˙W20 | 20 | 1.02e+10 | 39.1 | 329 | 57.3 | 0.299 |
| AD7˙W25 | 25 | 1.26e+10 | 39.4 | 191 | 56.6 | 0.25 |
| AD7˙W27 | 27 | 1.04e+10 | 39.5 | 423 | 56.3 | 0.467 |
| AD7˙W30 | 30 | 9.57e+09 | 39.7 | 268 | 58.3 | 0.71 |
| AD7˙W35 | 35 | 1.52e+10 | 39.9 | 233 | 55.4 | 0.703 |
| AD7˙W40 | 40 | 9.54e+09 | 39.8 | 146 | 54.4 | 0.641 |
| AD7˙W45 | 45 | 9.09e+09 | 39.9 | 23.4 | 55.7 | 0.419 |

Table S4: **Anaerobic digester time series samples.**

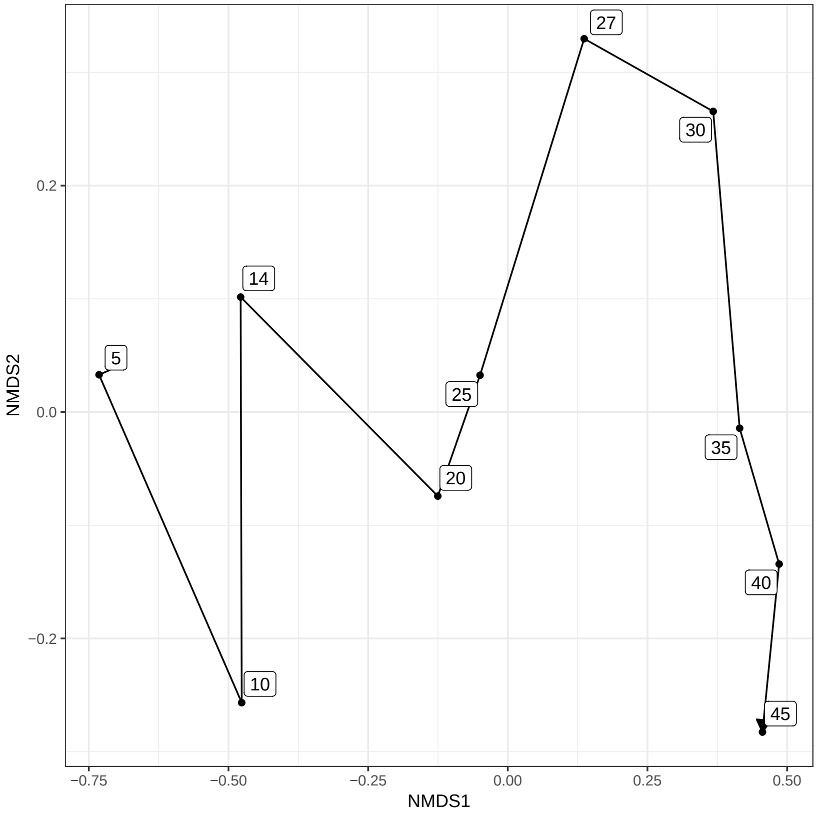


Figure S2: **NMDS plot of reactor community structure.** Bin coverages were normalised by per sample sequencing depth and Bray-Curtis distances calculated prior to NMDS, labels indicate sampling week. The sampling week had the strongest association with community structure ($R^2 = 0.46$, $p = 0.001$) followed by $H_2S$ concentration ($R^2 = 0.17$, $p = 0.009$) and $O_2$ ($R^2 = 0.14$, $p = 0.014$) based on PERMANOVA with Bray-Curtis distances (Adonis function in vegan library R). The other operating conditions, temperature and $CH_4$, were not significantly associated.

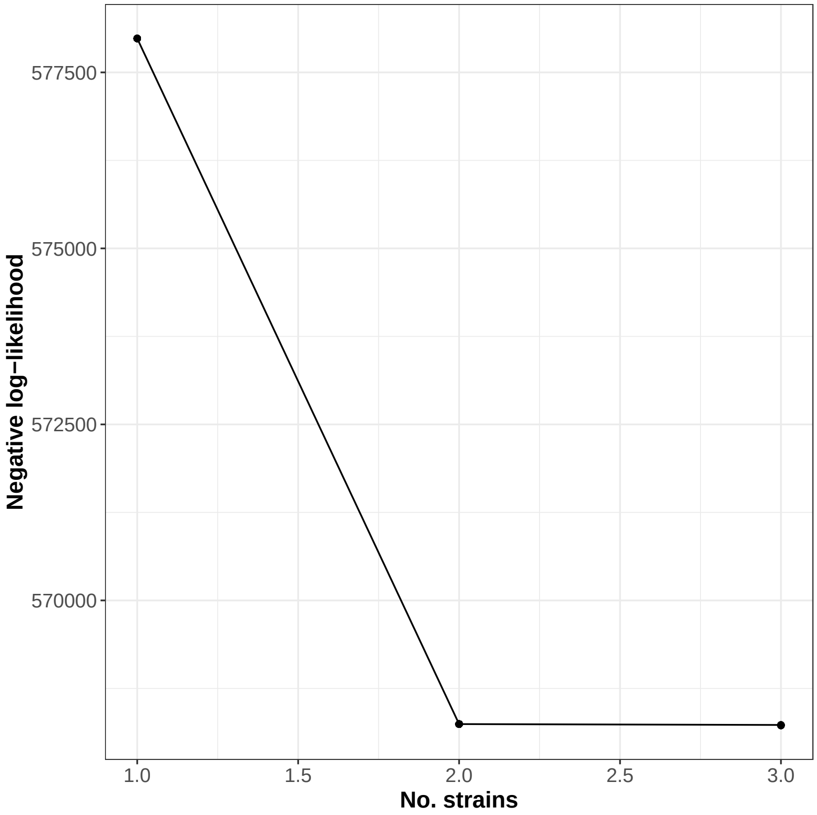


Figure S3: **Negative log-likelihood of Nanopore haplotype fits as a function of strain number for COG0532 from Bin_72 of the AD time series data set.** The hybrid EM algorithm for Nanopore strain resolution defined in the Methods was applied to all 1,603 Nanopore reads mapping to this SCG and the negative log-likelihood computed for ten replicates at each strain number. The algorithm was run for up to 4 strains but degenerate haplotypes are collapsed and in practice no more than three strains were ever observed in this case.

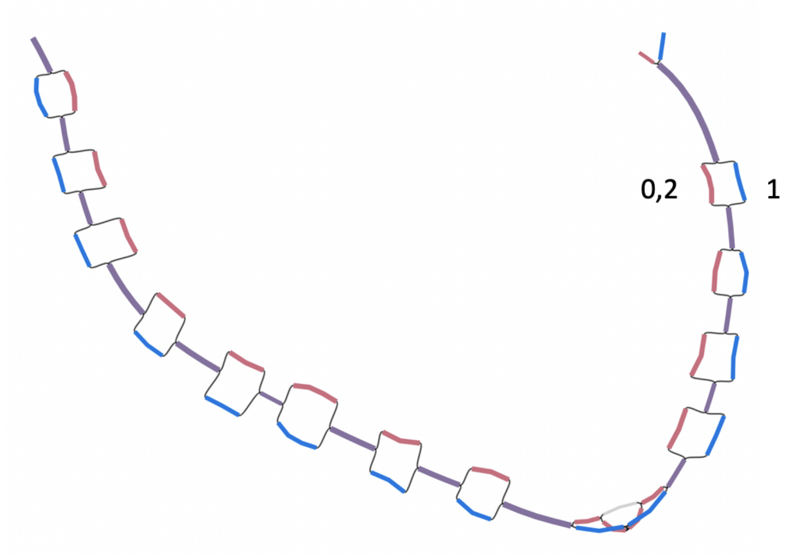


Figure S4: **Simplified variant graph for COG0532 from Bin_72 of the AD time series**. This is a bandage plot of the high-resolution subgraph extracted for COG0532 post-simplification. The colours indicate which unitig is present in which strain (0,1 and 2). In this case strains 0 and 2 are identical for this gene.

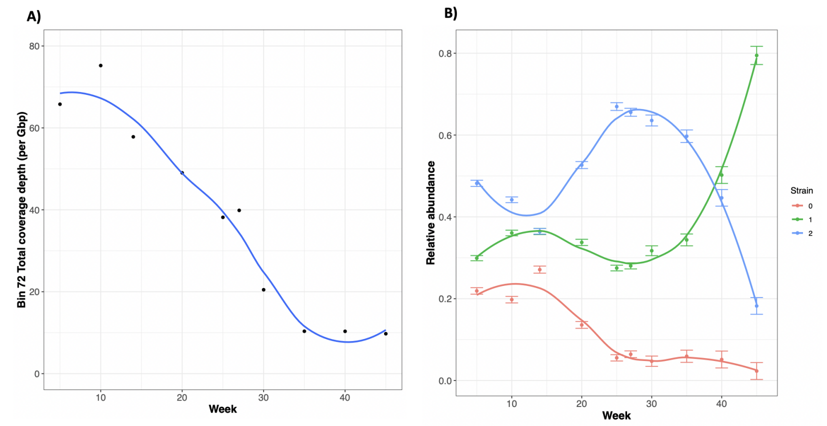


Figure S5: **Time series of A) total MAG coverage depth and B) STRONG strain abundances from Bin_72 of the AD time series**. A) Coverage depth of Bin_72 normalised by Gbp of sequence in each sample. This MAG decreased significantly in abundance over time (Pearson's correlation on log coverage $r = -7.9$ Benjamini-Hochberg adjusted $p = 4.9e - 05$). B) Strain relative proportions as calculated by BayesPaths with uncertainties as twice the standard deviation in the variational Bayesian prediction. Curves are LOESS smoothings of data points. Strain proportions did not change significantly (permutation multivariate ANOVA $R^2 = 0.35$ adjusted $p = 0.089$).

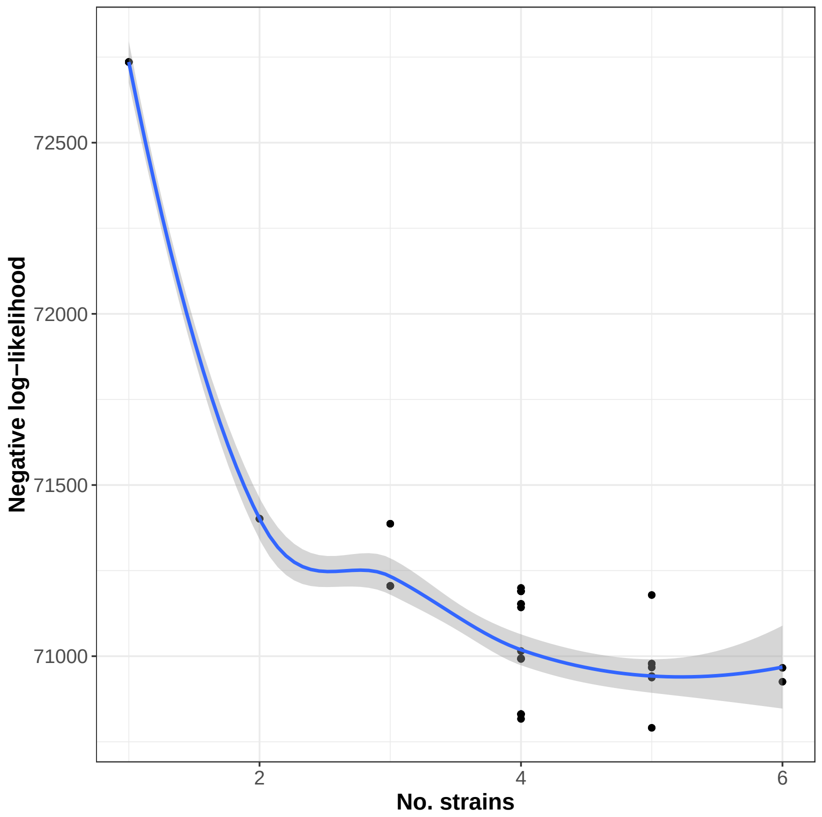


Figure S6: **Negative log-likelihood of Nanopore haplotype fits as a function of strain number for COG0072 from Bin_846 of the AD time series.** The hybrid EM algorithm for Nanopore strain resolution defined in the Methods was applied to the 194 Nanopore reads mapping to this SCG and the negative log-likelihood computed for ten replicates at each strain number. The algorithm was run for up to 6 strains.

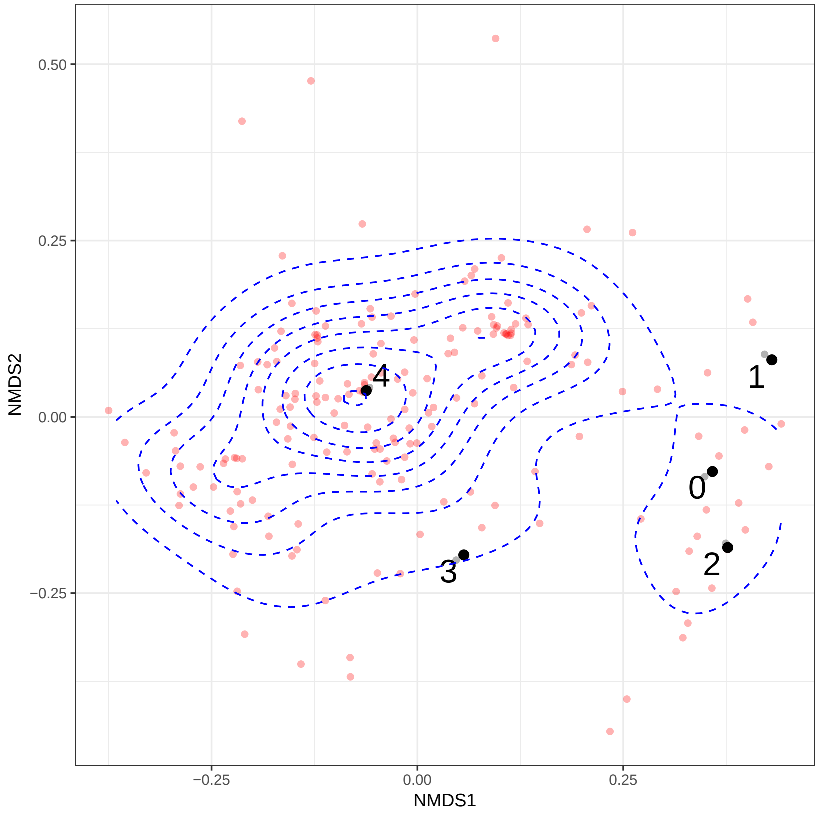


Figure S7: **Comparison of Nanopore reads to STRONG prediction for COG0072 from Bin_846.** Non-metric multidimensional scaling of Nanopore reads that mapped to COG0072 from Bin_846 of the anaerobic digester time series (red) together with the five haplotypes reconstructed from short reads by STRONG (black 0, 1, 2, 3 and 4). Distances were calculated as fractional Hamming distances on short read variant positions (see Methods). Blue dashed lines indicate read density contours. In this sample (Week 27) the short read haplotypes were predicted to have relative abundances $\rho_0 = 0.043 \pm 0.012$, $\rho_1 = 0.024 \pm 0.014$, $\rho_2 = 0.13 \pm 0.017$, $\rho_3 = 0.059 \pm 0.018$ and $\rho_4 = 0.74 \pm 0.027$.

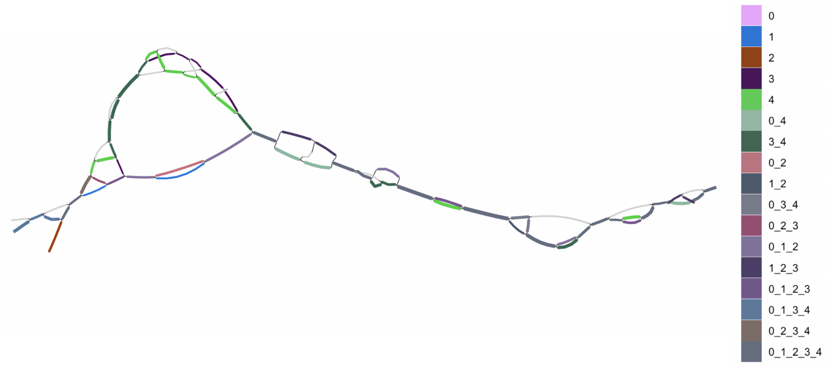


Figure S8: **Simplified variant graph for COG0072 from Bin_846 of the AD time series**. This is a bandage plot of the high-resolution subgraph extracted for COG0072 post-simplification. The colours indicate which unitig is present in which strain (see legend).

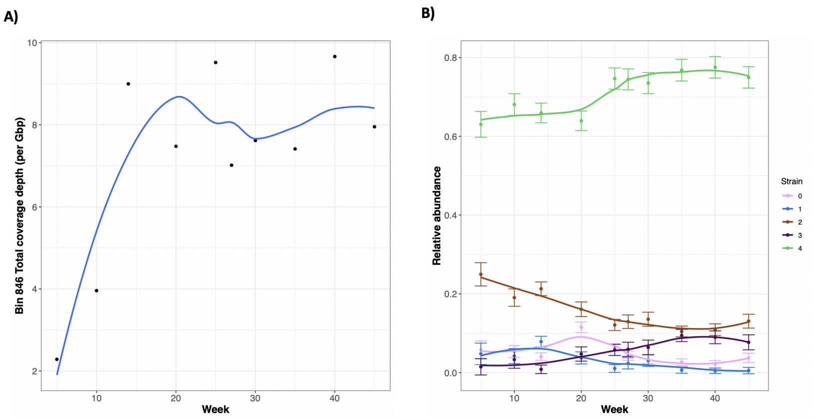


Figure S9: **Time series of A) total MAG coverage depth and B) STRONG strain abundances from Bin_846 of the AD time series**. A) Coverage depth of Bin_846 normalised by Gbp of sequence in each sample. This MAG increased marginally in abundance over time (Pearson's correlation on log coverage $r = 2.6$ Benjamini-Hochberg adjusted $p = 0.071$). B) Strain relative proportions as calculated by BayesPaths with uncertainties as twice the standard deviation in the variational Bayesian prediction. Curves are LOESS smoothings of data points. Strain proportions did change significantly (permutation multivariate ANOVA $R^2 = 0.72$ adjusted $p = 0.011$).

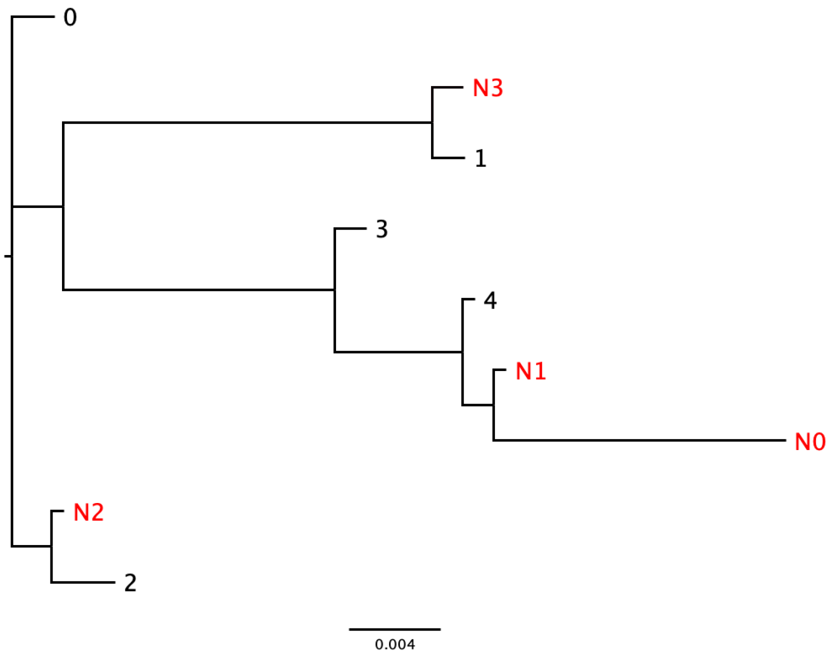


Figure S10: **Comparison of Nanopore and STRONG strain haplotypes for COG0072 from Bin_846 of the AD time series**. The four Nanopore haplotypes from the optimum run of the EM algorithm are shown in red and 5 STRONG haplotypes in black. The Nanopore haplotypes had relative abundance $\rho_0 = 0.056$, $\rho_1 = 0.820$, $\rho_2 = 0.091$ and $\rho_3 = 0.035$. N0 matched best to 4 with 98.8% nucleotide identity, N1 to 4 with 99.9%, N2 to 0 with 99.7%, N3 to 1 with 99.8%.
